# Structural divergence of G6PD variants shapes red blood cell storage and transfusion performance

**DOI:** 10.1101/2025.11.21.689741

**Authors:** Aaron V. Issaian, Shaun Bevers, Erin Marsh, Matthew S. Karafin, Roman R. Schlimgen, Julie A Reisz, Ariel Hay, Gregory R. Keele, Monika Dzieciatkowska, Francesca I. Cendali, Zachary B. Haiman, Alicia M. Key, Travis Nemkov, Daniel Stephenson, Amy L Moore, Mitasha S. Palha, Eric A. Legenzov, Derek R. Lamb, Xutao Deng, Mars Stone, Kirk C. Hansen, Steve Kleinman, Philip J. Norris, Michael P. Busch, Francesca Vallese, Erik W. Hartwick, James Jung, Bernhard O. Palsson, Steven L. Spitalnik, Joseph P.Y. Kao, Nareg H. Roubinian, Grier P Page, Elan Z. Eisenmesser, Lucio Luzzatto, James C Zimring, Paul W. Buehler, John Janetzko, Angelo D’Alessandro

## Abstract

Glucose-6-phosphate dehydroge-nase (G6PD) deficiency is the most common enzymopathy, yet how donor alleles shape red blood cell (RBC) storage, transfusion performance, and recipient physiology remains unresolved. Across 13,091 donors, a recalled cohort, donor–recipient data, humanized mice, and a prospective sickle cell disease cohort, variants reduced G6PD abundance, constrained pentose phosphate pathway flux and redox metabolism, and accelerated age-dependent functional decay. Murine A^−^ RBCs recovered poorly; A^−^ and Mediterranean RBCs impaired renal oxygenation, especially the latter, which caused greater plasma hemolysis. In humans, A^−^ units yielded smaller hemoglobin increments, while deficient units reduced allogeneic ^51^Cr post-transfusion recovery and four-week survival by 8 and 12-percentage-points. NADP⁺-bound cryo-EM resolved mechanisms: remodeling of the catalytic NADP⁺-binding region and dimer interface underlies A^−^ kinetic fragility, whereas Mediterranean S188F exhibits temperature-dependent oligomerization and conformational instability. These findings define an allelic series linking genotype to molecular instability, RBC quality, transfusion performance, and recipient physiology.

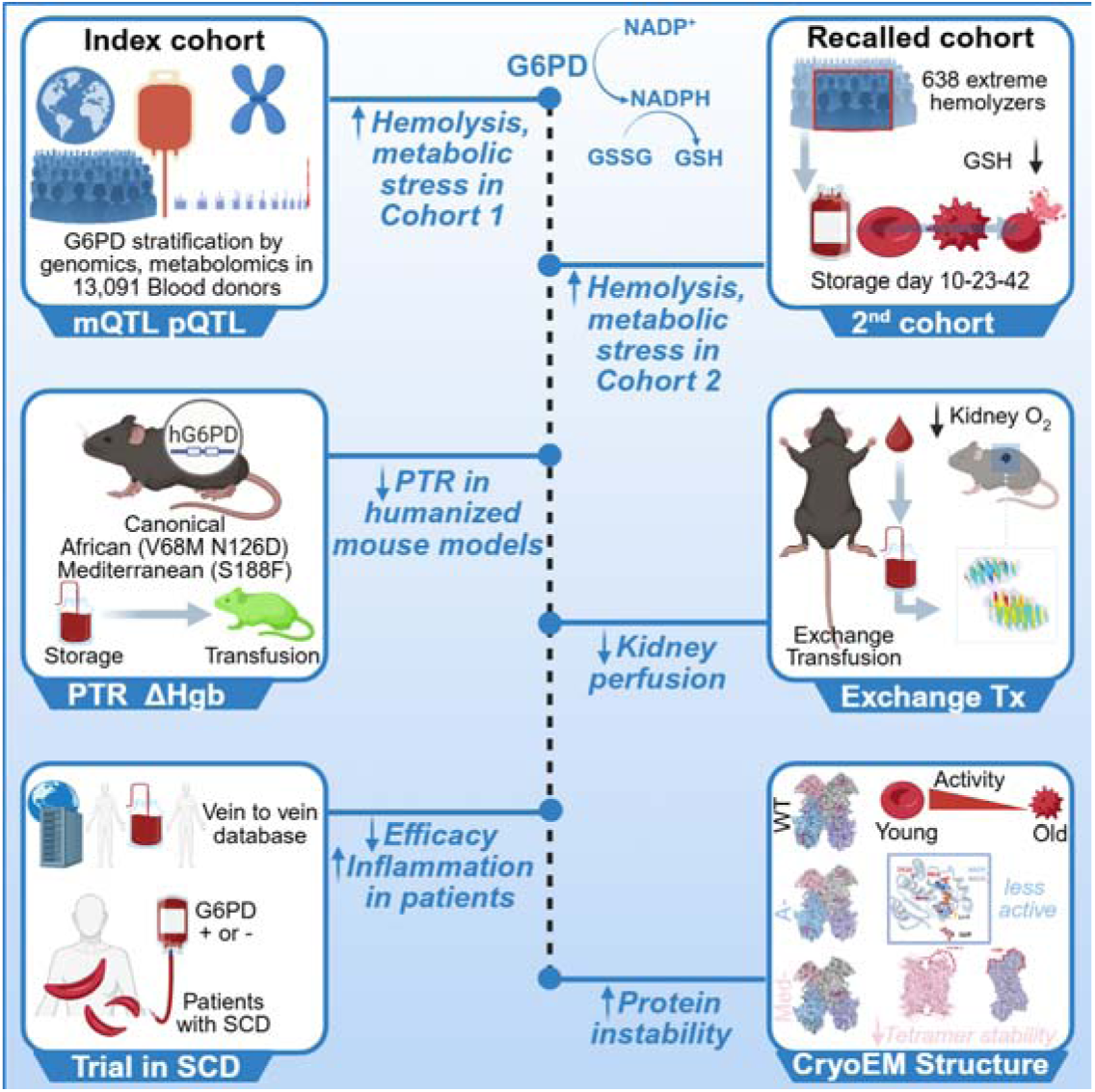

## INTRODUCTION

Glucose-6-phosphate dehydrogenase (G6PD) deficiency, the most common human enzymopathy, affects ∼500 million people worldwide^1^ and comprises >230 clinically relevant X-linked variants with differing catalytic activity, stability, and hemolytic risk.^2^ Red Blood Cells (RBCs) are especially vulnerable to this condition, because the oxidative pentose phosphate pathway (PPP) is their sole source of NADPH to fuel antioxidant systems, while their lack of nuclei and ribosomes precludes replenishment of unstable enzyme.^3,4^ Many disease-causing mutations cluster near the NADP⁺ binding site and the dimer interface, compromising thermodynamic stability and accelerating inactivation under oxidative load.^5–8^ These constraints may be amplified during refrigerated storage in the blood bank, when oxidative and membrane damage accumulate, G6PD activity declines even in leukoreduced units,^9,10^ and failure to activate the PPP may prime RBCs for post-transfusion clearance.^11–14^

Despite this biological rationale, donor G6PD status rarely informs blood inventory. Routine screening is not performed in the United States, and international policies vary; for example, NHS Blood and Transplant defers donors with known deficiency.^15,16^ Prevalence also varies with geography and donor selection: G6PD deficiency was identified in ∼0.3% of unselected RBC units at one metropolitan US service, ∼13% of male donors of African descent,^17,18^ a group where the burden of transfusion-dependent diseases is also highest.^19^ Indeed, donor G6PD-deficient status was confirmed in 12.3% of C−/E− antigen-matched units used for exchange transfusion in sickle cell disease (SCD).^17,20,21^ Among 3,004 Italian donors, 1.1% had reduced activity and nearly half of the deficient alleles were severe class II variants, including Mediterranean S188F;^22^ a 2026 survey of 110 Italian services found that only 19% accepted individuals with known deficiency for whole-blood donation.^23^ Thus, G6PD-deficient units enter the blood supply, including inventories serving chronically transfused populations, but their management remains inconsistent.

Evidence suggests impaired storage or transfusion performance, yet it does not establish a uniform effect across molecular variants. Genome-wide association analysis identified rs1050828 (V68M) as the leading G6PD signal for end-of-storage hemolysis after oxidant challenge,^24^ and targeted analysis linked this variant -which marks the African A^−^ haplotype when in cis with the N126D allele - to smaller post-transfusion hemoglobin increments.^25^ An autologous ^51^Cr study, the FDA-accepted reference for RBC recovery, found lower post-transfusion recovery (PTR) after 40–42 days of storage among G6PD-deficient donors.^17^ However, nine of ten deficient units were African A^−^, while the single Mediterranean unit had the lowest G6PD activity, yet comparatively preserved normal recovery. Concordantly, Mediterranean RBCs may appear preserved during refrigeration but hemolyze after rewarming to 37°C and oxidant challenge.^26^ In SCD, Sagiv et al. inferred reduced allogeneic donor-cell persistence from bulk HbA kinetics across serial, multi-unit transfusions,^19^ without directly tracking an individual unit. These observations implicate G6PD deficiency but leave unresolved which variants compromise stored RBCs, through what mechanisms, and whether these defects alter allogeneic persistence and recipient physiology.

Here, we addressed these gaps using complementary human and murine models together with structural, biochemical, and biophysical studies. We analyzed 362 G6PD SNPs in 13,091 blood donors enrolled in the Recipient Epidemiology and Donor Evaluation Study (REDS) RBC-Omics,^27^ with multi-omics data from 643 recalled donors and linked donor–recipient outcomes. Preclinically, we developed matched humanized knock-in models expressing canonical, complete African A^−^, or Mediterranean G6PD within the same murine regulatory context. These extend previous approaches focused either on humanized S188F G6PD, which showed no clear PTR deficit,^28^ or on V68M alone on a murine G6PD background,^29^ which showed poorer storage quality but does not reproduce the complete African A^−^ haplotype because it lacks the co-inherited N126D variant. We measured storage phenotypes and PTR and, after controlled hemorrhage without shock followed by serial exchange transfusion - a model relevant to hemorrhagic replacement and aspects of exchange transfusion in SCD^30^ - measured renal oxygenation in vivo.^31^ We also directly tracked a single ^51^Cr-labeled allogeneic unit in an exploratory prospective cohort of patients with SCD, combining the direct recovery measurement of Francis et al.^17^ with the clinical setting examined indirectly by Sagiv et al.^19^ Finally, we coupled in vivo RBC aging and biochemical assays with four near-atomic-resolution, NADP⁺-bound cryo-EM structures: sequence-intact canonical, African A^−^, and Mediterranean tetramers and a Mediterranean dimer. To our knowledge, these include the first NADP⁺-bound tetramer of sequence-intact WT human G6PD and the first structures of the complete African A− and Mediterranean variants, revealing distinct relationships among catalytic activity, oligomeric stability, and time-dependent functional decay.

## RESULTS

### Common missense G6PD variants are prevalent in blood donors and associate with oxidative and osmotic, but not storage hemolysis

In the REDS RBC-Omics cohort, 13,091 RBC units collected at four US blood centers were stored for 42 days, the end of shelf life in most countries, and analyzed by high-throughput metabolomics; 12,757 donors were genotyped across 362 *G6PD* SNPs (**Figure 1A**). Fifteen missense variants were detected (**Figure 1B**). The most frequent, N126D (A⁺; AX-37209045 imputed to rs1050829), occurred in ∼5% of donors, with dosage interpreted according to the X-linked inheritance of *G6PD*. A⁺ retains approximately 80–100% of normal activity and is classified as non-deficient: historical WHO Class IV and 2024 WHO Class C (>60% activity, no hemolysis).^2^ V68M (rs1050828), the second most frequent variant, is known as Asahi when present alone and forms the African A^−^ haplotype in cis with N126D. Because every V68M-positive donor also carried N126D, we refer to these donors as A^−^ throughout, whereas 199 donors carried exclusively A^+^ alleles. A^−^ was Class III under the legacy 1966/1985 scheme (∼10–60% residual activity) and is Class B under the 2022/2024 scheme (<45% activity; acute, trigger-dependent hemolysis). Class I Loma Linda/Aachen (N363K); Class II Mediterranean (S188F), Viangchan (V291M), Ierapetra (P353S), and Canton/Agrigento (R459L); and Class III Seattle (D282H) each had a minor allele frequency below 0.2%, consistent with reduced eligibility or participation among donors carrying more severe variants.

**Figure 1.**
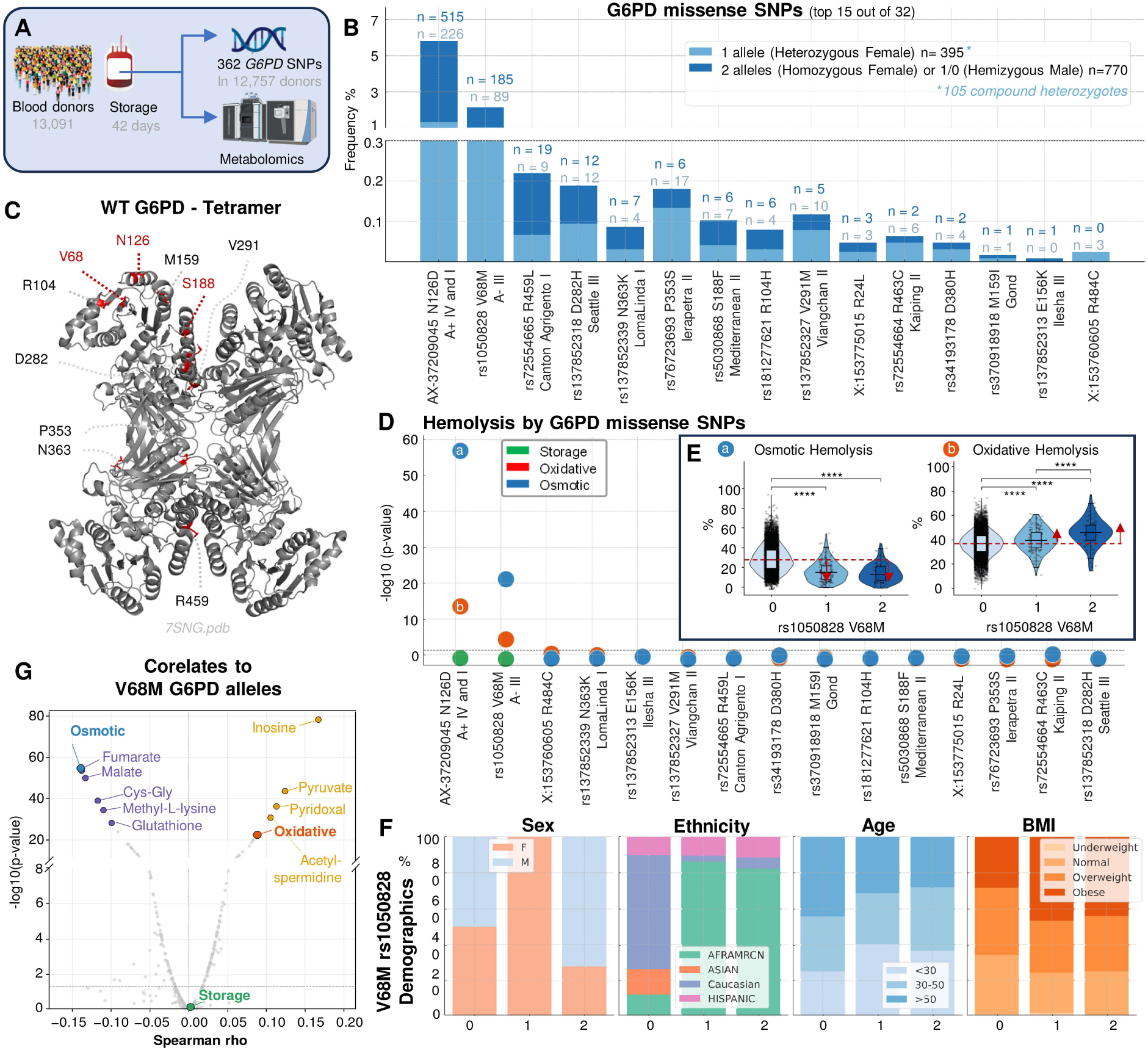
Common G6PD variants in blood donors associate with altered red blood cell metabolism and hemolytic propensity. REDS RBC-Omics design: 13,091 leukoreduced units stored to day 42 for metabolomics and 12,757 donors genotyped at 362 G6PD SNPs **(A)**. Frequencies of 15 missense variants were calculated accounting for X-linked inheritance; all V68M carriers also carried N126D in cis and are designated African A− **(B)**. Selected variants were mapped onto tetrameric human G6PD, including residues near catalytic and oligomerization interfaces **(C)**. Variant associations with storage, oxidative, and osmotic hemolysis are shown **(D)**. End-of-storage osmotic and oxidative hemolysis are stratified by V68M dosage **(E)**. Sex, self-reported ethnicity, age, and body mass index are shown by V68M dosage **(F)**. Correlations link V68M dosage to metabolites and hemolysis phenotypes **(G)**. See also Figure S1.

Although structures exist for canonical G6PD and selected pathogenic variants, including Canton and Viangchan,^5,6,32^ structures of full-length G6PD African A^−^ and Mediterranean S188F variants had not been determined. Mapping these residues onto the unliganded canonical tetramer (PDB:7SNG)^32^ showed that many cluster near the dimer interface or catalytic site, with potential effects on stability or catalysis, respectively (**Figure 1C**). Across variants, *G6PD* missense alleles associated with greater oxidative hemolysis and, more strongly, lower osmotic fragility, particularly for N126D (**Figure 1D**). The osmotic association persisted after excluding N126D-positive donors carrying V68M in cis, restricting the analysis to N126D without another detected clinically relevant *G6PD* variant. Dosage of rs1050828 showed graded effects on both hemolytic traits (**Figure 1E**). Donor stratification confirmed N126D prevalence (**Figure S1A**), enrichment of V68M among male donors of African descent, and enrichment of overweight and obesity among G6PD-deficient donors (**Figure 1F**), consistent with reported links to obesity and cardiovascular disease.^33^ Other missense variants, including S188F, Canton, and Seattle, showed distinct metabolic phenotypes and were enriched among donors of Asian or European ancestry (**Figure S1B–D**).

Uniform manifold approximation and projection (UMAP) of index-donor metabolomes showed the expected overlap between N126D and V68M, reflecting the occurrence of N126D independently as non-deficient A^+^ and in cis with V68M as African A^−^ (**Figure S1E**). Among rarer variants, Mediterranean S188F produced the most divergent metabolic phenotype and an apparent allele-dose response by linear discriminant analysis, consistent with its severity under both legacy Class II and current Class B designations (**Figure S1F**). At the end of storage, metabolites associated positively or negatively with V68M dosage (**Figures 1G** and **S1G**); several also correlated with spontaneous, oxidative, or osmotic hemolysis. Thus, *G6PD* genotype shapes the metabolic architecture and distinct dimensions of fragility in stored RBCs, with quantitatively and qualitatively different effects across variants.

### G6PD variation links cis regulation of protein abundance to redox metabolism

Metabolome-wide quantitative trait locus (mQTL) mapping linked *G6PD* variation most strongly to redox metabolites (**Figure 2A**), with ascorbate and 6-phosphogluconate providing representative regional signals at the *G6PD* locus (**Figure 2B–C**). A metabolite–SNP network identified rs1050828 (V68M; A^−^) as a major hotspot of metabolic variation (**Figure 2D**), enriched in NADPH-generating and -consuming pathways (**Figure 2E**), including glutathione homeostasis, the pentose phosphate pathway (PPP), glycolysis, and carnitine metabolism -the latter a hallmark of ferroptosis in stored RBCs.^34^ *G6PD* mQTLs also involved carboxylic acids such as fumarate and malate, which are linked to alternative NAD(P)H homeostasis^17,29^ through cytosolic Krebs-cycle enzyme isoforms in mature RBCs.^35,36^ V68M dosage was enriched among donors of African genetic ancestry (**Figure 2F**); however, ancestry-stratified analysis of glutathione disulfide (GSSG) retained a prominent chromosome-X *G6PD* signal, linking *G6PD* variation to RBC redox homeostasis independently of broad population structure (**Figure 2G**).

**Figure 2.**
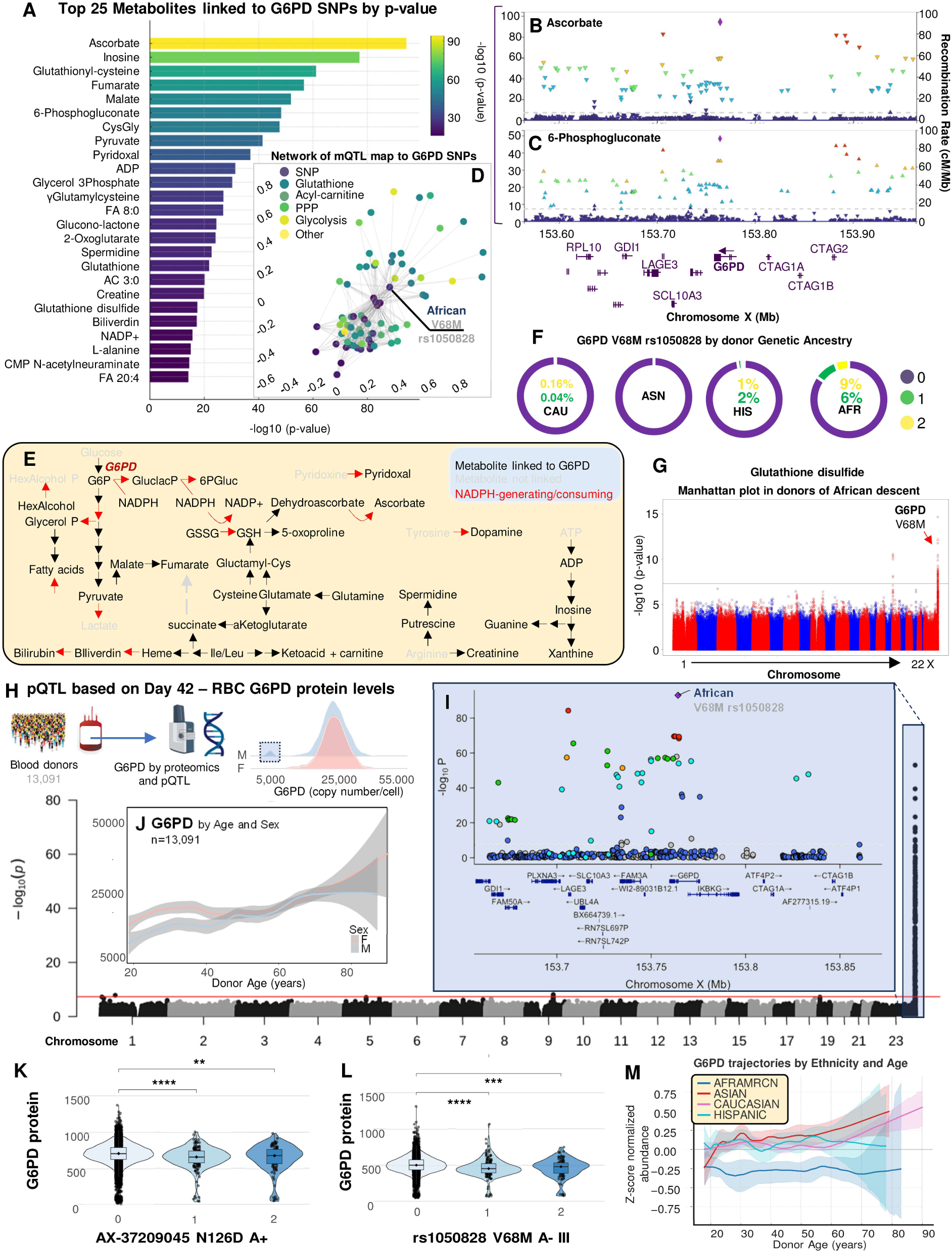
G6PD genetic variation shapes redox metabolism and G6PD protein abundance in human red blood cells. Metabolite quantitative trait locus (mQTL) results include ranked associations, regional signals for ascorbate and 6-phosphogluconate, a V68M-centered SNP-metabolite network, and pathways involving NADPH metabolism, the pentose phosphate pathway, glutathione, and glycolysis **(A–E)**. V68M enrichment by donor genetic ancestry and ancestry-stratified GWAS of glutathione disulfide are shown **(F–G)**. Day-42 G6PD protein quantification, protein QTL design, and the regional cis-pQTL signal are shown **(H–I)**. G6PD abundance is presented by donor age and sex, N126D dosage, V68M dosage, and age and genetic ancestry **(J–M)**. See also Figures S1 and S2.

Targeted proteomics of the same 13,091 REDS RBC-Omics units revealed sex-dependent G6PD copy number per cell and a distinct subgroup of male donors with very low protein abundance (**Figure 2H**). Protein QTL mapping identified a cis signal at *G6PD* (**Figure 2I**) and trans signals on chromosomes 9 and 19 near *HABP4* and *PLIN4*, both exceeding the genome-wide significance threshold of 5×10^-8^ (**Figure S1H–K**). Age and sex explained additional variance: females had higher G6PD abundance before age 51, the average age of menopause in the United States, and abundance increased with donor age (**Figure 2J**). Although it contrasts with previously-reported age-dependent declines,^37^ this pattern accords with the population bias introduced by “healthy-donor effects”,^38^ whereby older volunteers who remain eligible to donate constitute a selected population. Allele-dosage analyses distinguished the effects of A^+^ (N126D alone) from those of complete A^−^ (V68M in cis with N126D) on G6PD abundance (**Figure 2K–L**). Interactions between age and A^−^ further modulated abundance: the increase observed in noncarriers was absent among donors of African descent carrying V68M (**Figures 2M and S1I**). Thus, the *G6PD* locus exerts coordinated cis control of protein abundance and redox-focused metabolic remodeling in stored RBCs.

### Recalled-donor multi-omics and constraint-based modeling link genotype to flux remodeling

Of 13,091 index donors, 643 ranking above the 95th or below the 5th percentile for day-42 storage, oxidative, or osmotic hemolysis were recalled to donate a second unit. *G6PD* genotypes were available for 638, including 11 hemizygous A^−^ males and 7 heterozygous A^−^ females (**Figure S2A**); frequencies of the 15 principal missense variants resembled those in the index cohort (**Figure S2A–B**), despite the reported association of A^−^ with hemolytic traits^24^ and the broader associations identified here. Combined-omics linear discriminant analysis of units profiled by proteomics, metabolomics, and lipidomics at storage days 10, 23, and 42 separated samples primarily by storage duration and secondarily by rs1050828 dosage, with clinical, complete-blood-count, and multi-omic contributors defining this separation (**Figure S2C–D**). G6PD abundance decreased with V68M dosage throughout storage and was the strongest negative correlate of allele dosage across all omic features (**Figure S2E**). The N126-containing peptide supported this total-protein association, while 6-phosphogluconate and glutathione declined with storage and allele dosage (**Figure S2F–G**). As in the index cohort, rs1050828 dosage was associated with greater end-of-storage oxidative hemolysis but lower osmotic hemolysis (**Figure S2H**). In this cohort selected from the extremes of the hemolysis distributions, the sex difference observed in the index cohort was attenuated, whereas G6PD abundance declined with age (**Figure S2I**), consistent with previous reports.^37^

To infer the metabolic consequences of A^−^-associated proteomic changes, indexed by V68M dosage, we integrated longitudinal recalled-donor proteomics into proteome-constrained models, using enzyme abundance to constrain reaction capacities and flux variability analysis to estimate maximum feasible fluxes and effective ranges. The dependence of predicted flux on cognate-enzyme abundance varied widely across reactions (**Figure S2J**). G6PD abundance strongly predicted flux through G6PDH2 (ρ = 0.9652; **Figure S2J–K**); estimated ranges in heterozygous females and hemizygous males carrying V68M were approximately 50% and 10% of those in noncarriers, respectively. By contrast, downstream GND flux correlated weakly with PGD abundance (ρ = 0.0813) but paralleled the genotype-stratified G6PDH2 profile, consistent with upstream substrate supply limitation (**Figure S2K**).

Among genotype-associated flux differences meeting *P* < 2×10^-4^, reduced oxidative PPP flux coincided with increased ether-lipid flux (**Figure S2L**), including acyltransferase and phospholipase A₂ reactions in ether-linked phosphatidylcholine metabolism. Carriers of V68M also showed increased predicted glucose- and galactose-mutarotase fluxes (GLCM and GALM) and altered phosphoglucomutase activity. Integrated enrichment implicated PPP/glutathione and glycolytic metabolism; proteasomal and other protein-degradation pathways; autophagy; phospholipase D and mTOR signaling; ubiquitylation; and necroptosis (**Figure S2M**). Thus, A^−^-associated loss of G6PD abundance constrains oxidative PPP capacity and redistributes feasible flux toward sugar-isomerization and ether-lipid pathways.

Additional peptide-level analyses traced N126D, showed lower total G6PD throughout storage in N126D homozygotes/hemizygotes, and identified storage-dependent cysteine redox modifications and N-deamidation, together with evolving V68-and S188-diagnostic peptides consistent with the mQTL results (**Figures S2F and S3A–E**).

### Mouse models with humanized G6PD alleles recapitulate storage-dependent lesions and flux shifts

Previous transfusion models either introduced V68M alone by CRISPR-Cas9 on a C57BL/6J background^29^ or expressed S188F in a humanized, Cre-switchable strain retaining murine exons 1–2 along with a 13-bp scar between the promoter and start codon.^28^ The former does not reproduce African A^−^, in which V68M occurs in cis with N126D, and neither model directly compared complete A^−^, Mediterranean, and canonical human alleles at the endogenous locus. We therefore studied matched strains expressing non-deficient canonical, A^−^ (V68M/N126D), or Mediterranean (S188F) G6PD. Human *G6PD* exons 3–13 replaced the corresponding murine sequence without the scar, whereas murine exon 2 -retained to avoid disrupting overlapping IKKγ -and the endogenous murine regulatory context remained intact (**Figure 3A**).

**Figure 3.**
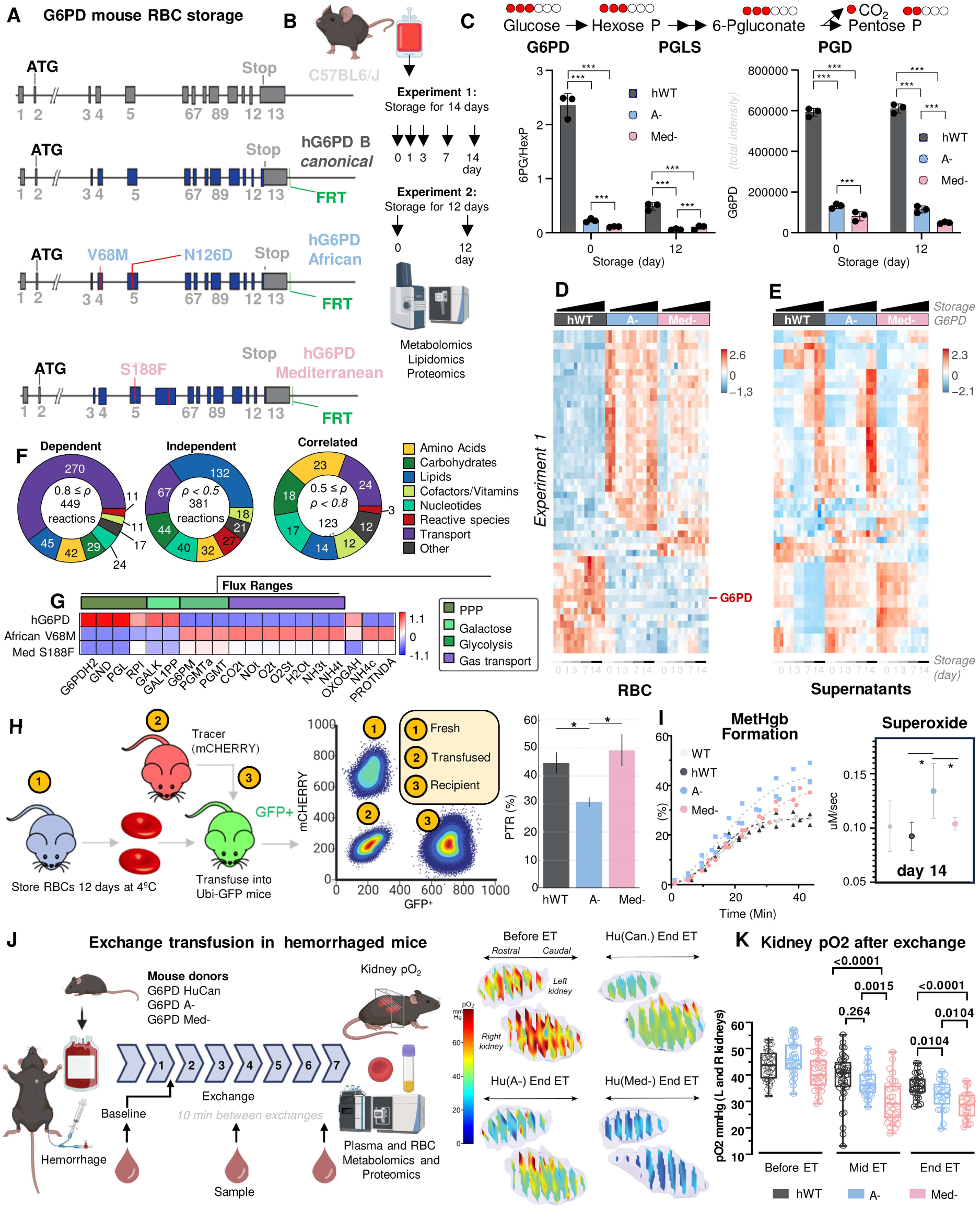
Humanized G6PD-deficient mice recapitulate storage-dependent metabolic dysfunction and impaired transfusion efficacy. Humanized loci encode canonical, African A− (V68M/N126D), or Mediterranean (S188F) G6PD within the endogenous murine regulatory context **(A)**. Complementary 4°C storage experiments sampled days 0, 1, 3, 7, and 14 or days 0 and 12 for metabolomics, lipidomics, and proteomics **(B)**. [1,2,3-^13^C_3_]glucose tracing and G6PD/PGD protein abundance show impaired oxidative pentose phosphate pathway (PPP) activity in deficient RBCs **(C)**. Storage- and genotype-dependent metabolomic and proteomic profiles are shown **(D–E)**. Proteome-constrained modeling classifies reactions by protein-abundance dependence and compares genotype-specific flux ranges **(F–G)**. Twelve-day-stored test RBCs were co-transfused with fresh mCherry⁺ tracers into GFP⁺ recipients; tracer-normalized 24h post-transfusion recovery (PTR) is shown **(H)**. Methemoglobin formation and day-14 superoxide generation are shown **(I)**. The controlled-hemorrhage model, serial exchange-transfusion with 7-day-stored RBCs and representative functional-MRI renal oxygenation maps are shown **(J)**. Renal pO₂ before, midway through, and after exchange transfusion is quantified **(K)**. See also Figures S3 and S4.

Donor RBCs were stored with longitudinal sampling; deep multi-omics focused on days 0,1,3,7 and 14 (experiment 1) or 0 and 12 (experiment 2 -**Figure 3B**). Tracing with 1,2,3-^13^C_3_-glucose showed reduced oxidative PPP flux in both deficient strains and a storage-dependent decline in canonical RBCs (**Figure 3C**), consistent with human data.^9,10^ Across two independent experiments, G6PD abundance was lower in both deficient strains, whereas PGD, the downstream NADPH-generating PPP enzyme, was selectively increased in A^−^ RBCs (**Figures 3D–E and S3G–H**). Multi-omic trajectories revealed genotype-by-storage interactions: PPP metabolites, free fatty acids, and oxidized lipids changed in proportion to deficiency, whereas conjugated bile-acid changes predominated in Mediterranean RBCs (**Figures 3D–E and S3G–H**). Stored A^−^ RBCs also showed higher levels of NRF2-regulated proteins,^39^ including NQO1 and PGD, consistent with compensatory programming during erythroid maturation rather than transcriptional adaptation in mature RBCs (**Figure 3E**). Supernatant heme increased during storage, with no significant pairwise differences among strains (**Figure S3F**), compatible with the lack of association between G6PD deficiency and storage hemolysis in REDS.^24^

Flux variability analysis of proteome-constrained models from all three strains estimated maximum feasible fluxes and effective ranges for 953 enzyme-catalyzed reactions carrying non-zero flux (**Figures 3F–G and S4**). Spearman correlations between maximum flux and cognate-enzyme abundance classified 381 reactions as abundance-independent (ρ < 0.5), 123 as abundance-correlated (0.5 ≤ ρ < 0.8), and 449 as abundance-dependent (ρ ≥ 0.8) (**Figure 3F**). Genotype-associated differences altered effective ranges in carbohydrate metabolism and membrane transport; functional grouping identified abundance-dependent water- and gas-transport reactions that connected aquaporin abundance to osmotic fragility (**Figures 3G and S4**). The G6PD reaction was abundance-dependent (ρ = 0.861), whereas the downstream NADPH-producing PGD reaction was not (ρ = 0.0548); nevertheless, modeled oxidative PPP capacity contracted markedly in both deficient strains, identifying G6PD abundance as the principal proteomic constraint on NADPH regeneration (**Figures 3G and S4**). Models also predicted altered hexose-phosphate mutarotase and mutase capacities, consistent with strong abundance dependence of PGMT and PGMTa (ρ = 0.9784; **Figure S4**). Thus, allele-dependent protein abundance constrains PPP capacity while remodeling carbohydrate and membrane-transport fluxes.

### African A^−^ G6PD selectively lowers murine PTR

To test whether the concordant murine and human redox phenotypes affected transfusion efficacy, we measured post-transfusion recovery (PTR) in the humanized models. An initial storage time course showed limited genotype separation in donor-RBC omics at day 7 (**Figure 3D-E**), when canonical RBCs retained >90% PTR. We therefore used day 12 as a sensitized interval capable of revealing emerging PTR defects (**Figure S3G-H**), consistent with previous murine studies.^40^ RBCs stored for 12 days from canonical, A^−^, or Mediterranean mice were co-transfused with fresh mCherry⁺ reference RBCs into Ubi-GFP recipients; recovery of test cells at 24 h was normalized to that of the reference cells (**Figure 3H**). PTR was significantly lower for A^−^ RBCs than for either canonical or Mediterranean RBCs, demonstrating an allele-specific effect not simply proportional to conventional deficiency severity. Upon *in vitro* challenge with pro-oxidant primaquine orthoquinone, Methemoglobin formed faster and was reduced less efficiently in both deficient strains, most prominently in A^−^ (**Figure 3I**). At day 14, electron paramagnetic resonance detected greater superoxide production in A^−^ RBCs, (**Figure 3I**).

### Donor G6PD variants differentially impair renal oxygenation and increase recipient hemolysis markers after hemorrhage in mice

To assess tissue oxygen delivery before overt storage failure, we used the controlled-hemorrhage exchange-transfusion model of Baek et al.^30^ A defined blood volume was removed without inducing shock, followed by seven sequential exchanges with canonical, A^−^, or Mediterranean RBCs stored for 7 days (**Figure 3J**). At this interval, canonical RBCs retained >90% PTR, allowing physiological differences to be examined before the broader deterioration evident after 12 days. Avoiding shock minimizes systemic hemodynamic and inflammatory injury, while progressive RBC replacement reproduces a central procedural feature of exchange transfusion in SCD^41^ without modeling its underlying pathobiology. Because ex vivo perfusion studies show that renal oxygenation is sensitive to dysfunctional RBC O₂ kinetics,^31^ electron paramagnetic resonance imaging quantified renal pO₂ at baseline, mid-exchange, and after the seventh exchange (**Figures 3J–K**). Baseline pO₂ was comparable among groups. At mid-exchange, recipients of Mediterranean RBCs were significantly impaired, whereas A^−^ recipients had an intermediate phenotype not significantly different from canonical controls. After seven exchanges, renal pO₂ was reduced with both deficient RBC types, most markedly with Mediterranean RBCs (**Figure 3K**), demonstrating a donor-genotype effect on organ-level transfusion efficacy beyond circulating-cell recovery.

Recipient RBC and plasma multi-omics revealed distinct genotype-dependent responses (**Figure S4**). RBCs from A^−^ recipients diverged most from the other groups and showed lower G6PD abundance, 6-phosphogluconate, and reduced glutathione, consistent with persistence of a remodeled population with compromised PPP capacity and redox buffering. Conversely, plasma from Mediterranean recipients showed the greatest omic divergence and highest levels of hemolysis-associated cell-free Hba and Hbb-b1, LDH, and bilirubin, followed by A^−^ recipients (**Figure S4**). These patterns support relatively clearance-dominant A^−^ and plasma-hemolysis-dominant Mediterranean phenotypes, without defining the anatomical route or relative intra- and extravascular contributions to RBC loss. Across these complementary paradigms, preserved Mediterranean PTR after 12-day storage did not predict preserved renal oxygenation after 7-day serial exchange, disentangling RBC circulatory persistence from preserved function.

### The African A− G6PD haplotype is associated with reduced hemoglobin increments in humans

Having observed allele-specific post-transfusion phenotypes in mice, we asked whether this heterogeneity extended to humans. Previous studies identified rs1050828 as the principal G6PD signal for *in vitro* oxidative hemolysis,^24^ linked donor G6PD deficiency to smaller post-transfusion hemoglobin increments,^25^ and found lower autologous ^51^Cr recovery of deficient units after 40–42 days of storage.^17^ The latter cohort comprised nine African A^−^ donors and one Mediterranean donor, whose unit paradoxically retained 81.4% PTR despite having the lowest measured G6PD activity. Allogeneic evidence in SCD instead relied on ΔHbA kinetics across serial, multi-unit transfusions rather than direct tracking of an individual unit.^19^ Because these studies did not resolve recipient outcomes by allele, we tested whether transfusion efficacy tracked G6PD deficiency uniformly or selectively.

Allele-resolved analysis of the vein-to-vein database,^42^ comprising 4,464 single-unit transfusion events, showed a dose-dependent reduction in adjusted hemoglobin increment (ΔHb; post-minus pre-transfusion concentration) with donor rs1050828 (**Figures 4A–B**). Because every rs1050828-positive REDS donor also carried N126D in cis, this variant indexed the complete African A^−^ V68M/N126D haplotype. Adjusted ΔHb (95% CI) was 0.99 g/dL (0.97–1.01) for noncarrier units, 0.79 g/dL (0.52–1.07) at dosage one, and 0.63 g/dL (0.36–0.89) at dosage two - a 36% reduction relative to noncarriers (**Figure 4A**). The intronic variant rs743544, in linkage disequilibrium with A^−^, showed a concordant association: adjusted ΔHb fell from 0.95 g/dL (0.91–0.98) in noncarriers to 0.42 g/dL (0.00–0.86) at dosage two, a 56% reduction (**Figure 4A**). No comparable dose-dependent reduction was detected for Mediterranean S188F (rs5030868), but few informative events produced wide confidence intervals and precluded concluding equivalent efficacy. Thus, lower ΔHb tracked with African A^−^ and its linked intronic signal rather than uniformly across tested G6PD variants. We next directly measured allogeneic donor-cell recovery and survival in recipients with SCD.

**Figure 4.**
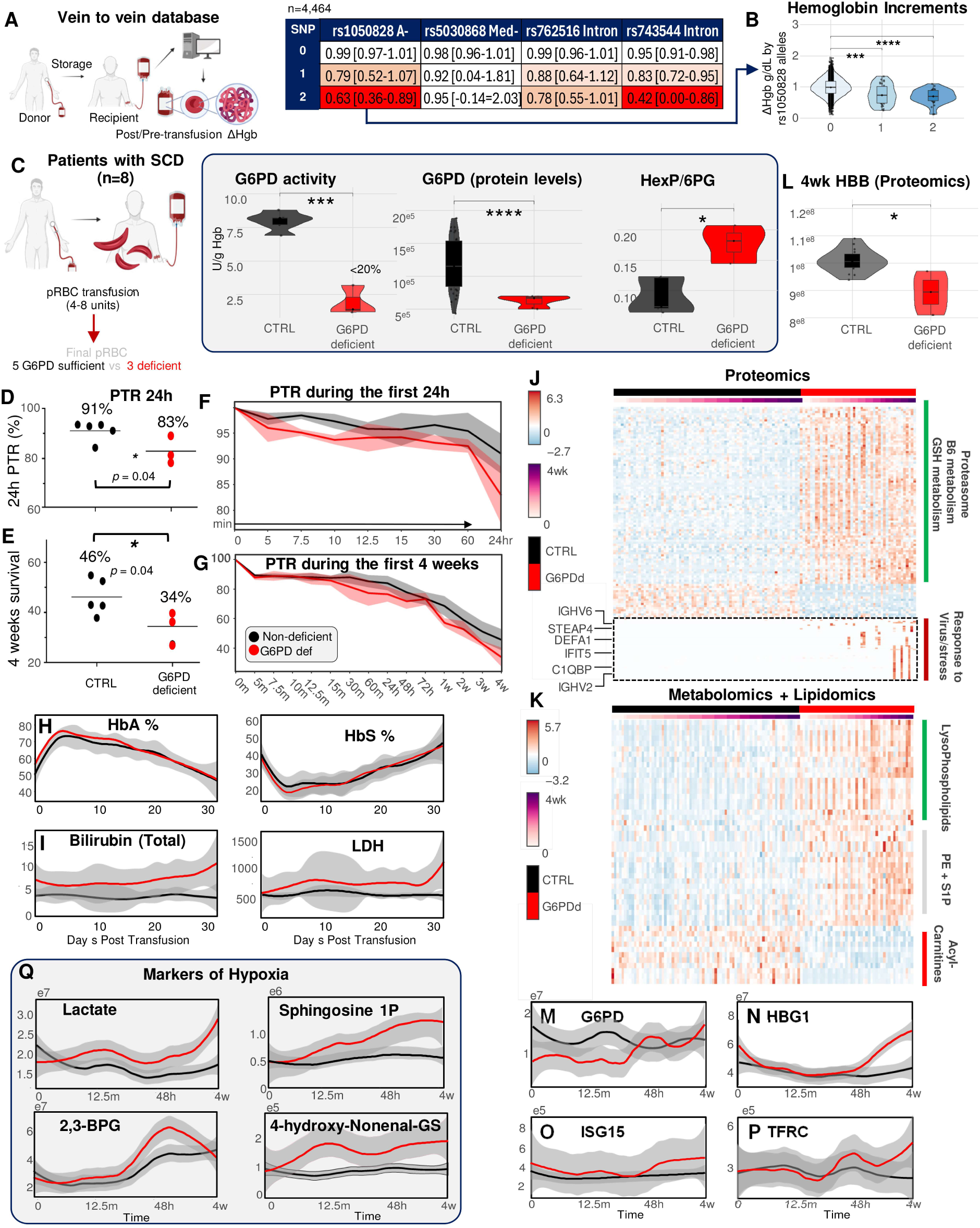
Donor G6PD deficiency reduces transfusion efficacy and promotes hemolytic and hypoxic responses in humans. In 4,464 single-unit transfusions, adjusted hemoglobin increments and confidence intervals are reported for selected donor G6PD variants **(A)**. Hemoglobin increments are stratified by V68M dosage **(B)**. Eight patients with sickle cell disease received 4–8 exchange units; all were G6PD-sufficient except the final ^51^Cr-labeled study unit, which was G6PD-sufficient (n = 5) or deficient (n = 3). Activity, G6PD protein, and the hexose phosphate/6-phosphogluconate ratio confirmed donor phenotype **(C)**. Unit-specific 24h PTR, first-24h recovery trajectories, four-week survival, and four-week trajectories are shown **(D–G)**. HbA/HbS and total bilirubin/LDH were followed longitudinally **(H–I)**. Recipient RBC proteomic and metabolomic/lipidomic profiles are shown **(J–K)**. Non-sickle HBB abundance at four weeks by proteomics is shown **(L)**. Longitudinal G6PD, HBG1, ISG15, and TFRC abundance is shown **(M–P)**. Lactate, sphingosine-1-phosphate, 2,3-bisphosphoglycerate, and 4-hydroxynonenal-glutathione trajectories are shown **(Q)**.

### Prospective direct tracking of allogeneic G6PD-deficient RBCs in SCD shows lower recovery and survival

Participants received 4–8 packed RBC units, all of which were G6PD-sufficient except the final, which was either G6PD-sufficient (n=5; median activity, 8.2 U/gHb; range, 6.9–8.8) or deficient (n=3; median, 1.4 U/gHb; <20% of normal). Only the final unit in each study arm was ^51^Cr-labeled. Omics data confirmed lower G6PD abundance and an elevated hexose phosphate-to-6-phosphogluconate ratio, independently confirming deficiency (**Figure 4C**). Whereas Francis et al. measured autologous ^51^Cr recovery in healthy donors after 40–42 days of storage,^17^ we followed an allogeneic unit in recipients with SCD. Unlike the bulk HbA kinetics used by Sagiv et al. across serial, multi-unit transfusions,^19^ direct labeling isolated the study unit from minutes through four weeks against a G6PD-sufficient exchange background. To our knowledge, this is the first prospective direct measurement of ^51^Cr-recovery and four-week survival for an allogeneic G6PD-deficient unit in SCD.

The prespecified primary endpoint, mean 24h ^51^Cr PTR, was 91.03% for sufficient and 82.91% for deficient units (difference, −8.12-percentage-points; two-sided 95% Welch CI, −19.33 to 3.09; exact one-sided Mann–Whitney *P* = 0.0357; **Figure 4D**), with separation emerging during the first 24h (**Figure 4E**). Mean four-week survival was 46.14% and 34.40%, respectively (difference, −11.75-percentage-points; two-sided 95% Welch CI, −24.47 to 0.97; exact one-sided Mann–Whitney *P* = 0.0357; **Figure 4F**), and longitudinal trajectories sustained this difference (**Figure 4G**).

Unadjusted HbA and HbS percentages, previously used to infer donor-cell persistence across multi-unit transfusions,^19^ did not clearly differ between groups (**Figure 4H**). These bulk fractions integrate all 4–8 units and cannot isolate the labeled unit. Total bilirubin and LDH trajectories were higher after deficient-unit transfusion, compatible with greater hemolytic activity, although these secondary observations are exploratory (**Figure 4I**).

Exploratory recipient-RBC multi-omics revealed group-associated proteomic, metabolomic, and lipidomic trajectories through four weeks (**Figures 4J–K**). Non-sickle HBB abundance was lower at four weeks after deficient-unit transfusion, consistent with reduced donor-RBC persistence (**Figure 4L**), while G6PD, HBG1, ISG15, and TFRC followed divergent trajectories (**Figures 4M–P**). The broader proteomic signature involved glutathione and proteasomal metabolism; complement and interferon responses,^43^ including C1QBP and IFIT5; iron homeostasis, including STEAP4; and immunoglobulin proteins IGHV2 and IGHV6 (**Figure 4J**). Metabolomic and lipidomic changes included higher lysophospholipids, phosphatidylethanolamines, and sphingosine-1-phosphate but lower acylcarnitines, which have been associated with kidney dysfunction^44^ (**Figure 4K**). Lactate, sphingosine-1-phosphate, and 2,3-bisphosphoglycerate -metabolites associated with hypoxic adaptation and oxygen delivery^45,46^ - and the lipid-oxidative-stress marker 4-hydroxynonenal– glutathione were also higher (**Figure 4Q**). These secondary findings provide hypothesis-generating evidence of sustained hypoxic, oxidative, and inflammatory remodeling, coupling unit-specific recovery and survival to recipient physiology.

### G6PD variants undergo distinct modes of functional decay during RBC aging

To determine whether progressive dysfunction during storage and circulation reflects age-dependent enzyme loss *in vivo*, we age-resolved RBCs by biotinylation in the humanized mice, thereby avoiding potential logistical and ethical constraints of labeling in human, while achieving greater precision than Percoll density fractionation.^47^ G6PD activity was highest in newly generated cells (**Figure 5A**): biotin-negative RBCs ≤6 days-old from A^−^ and Mediterranean mice retained approximately 95% and 60% activity, respectively, relative to the total RBC population from canonical mice (**Figure 5A**). Younger cells of all strains also had greater G6PD abundance and PPP activity (**Figure 5B**). These data demonstrate variant-dependent decay of G6PD abundance and activity during the RBC lifespan, consistent with reported half-lives of 62, 13, and <1 day for canonical, A^−^, and Mediterranean G6PD, respectively.^48^

**Figure 5.**
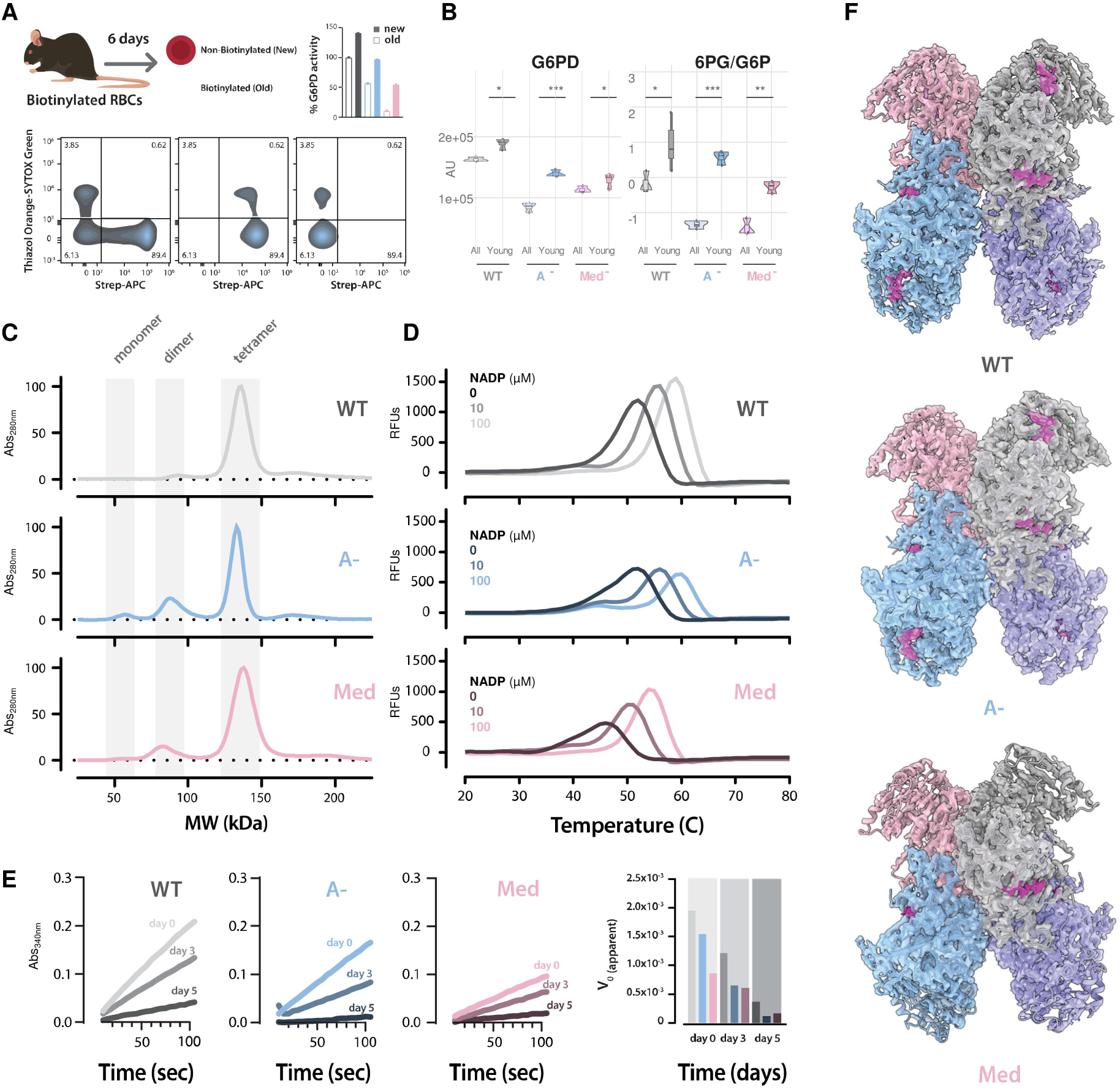
G6PD-deficient variants undergo distinct modes of functional decay during red blood cell aging. Six days after in vivo biotin labeling, newly generated and previously circulating RBCs were resolved for age-dependent G6PD activity **(A)**. G6PD abundance and PPP activity, indexed by the 6-phosphogluconate/hexose phosphate ratio, were compared in young and total RBC populations from canonical, A−, and Mediterranean mice **(B)**. Recombinant enzyme activity during incubation at 37°C reveals accelerated A− functional decay and lower Mediterranean baseline activity **(C)**. Single-molecule mass distributions show variant-dependent oligomeric states **(D)**. Thermal unfolding profiles were measured with increasing NADP⁺ **(E)**. NADP⁺-bound (magenta color) canonical, A−, and Mediterranean G6PD tetramers were resolved by cryo-EM **(F)**. See also Figures S5 and S6.

Existing structures have largely examined unliganded canonical tetramers, NADP⁺-bound dimers, or variants at the dimer–dimer interface, including Canton and Viangchan;^5–7,32^ neither Mediterranean G6PD nor the complete active-site-proximal A^−^ proteoform had been resolved. We therefore purified recombinant canonical, A^−^ (V68M/N126D), and Mediterranean (S188F) enzymes (**Figure S5A**). Diagnostic-peptide mass spectrometry confirmed sequence fidelity, while single-molecule mass distributions revealed variant-specific oligomeric states (**Figure 5C**). Measured *k*cat values of 323, 150, and 89 s^-1^, respectively (**Figure S5B**), agreed with reported values of approximately 275, 137, and 98 s^-1^, independently validating the freshly purified proteins.^15^ After one week, mass photometry showed that canonical G6PD remained predominantly tetrameric at 4°C, whereas incubation at 37°C shifted the population toward dimers (**Figure S5C**).

Fluorescence-based thermal unfolding further distinguished the variants: A^−^ remained closer to canonical, whereas Mediterranean G6PD transitioned at lower temperatures. Increasing NADP⁺ from 0 to 10 and 100 μM shifted transitions upward, including those of Mediterranean G6PD (**Figure 5D**), consistent with NADP⁺ stabilization of G6PD oligomers.^49^ DSC likewise showed Mediterranean transitions approximately 3.7°C below canonical and reduced tetramer-transition calorimetric enthalpy (126 versus 144 kcal mol^-1^). A^−^ transition temperatures remained near canonical, but its first-transition apparent van’t Hoff enthalpy was lower (172 versus 252 kcal mol^-1^), indicating altered apparent cooperativity rather than the pronounced destabilization reported for N126D/L323P.^50^

Fresh recombinant Mediterranean G6PD had ∼60% of canonical activity, whereas A^−^ retained >90% (**Figure 5E**). At 37°C, however, A^−^ lost activity faster than canonical G6PD and by day 5 fell below Mediterranean, revealing kinetic fragility despite near-normal initial activity (**Figure 5E**). Quantitative TMT crosslinking mass spectrometry likewise showed greater temperature-dependent intramolecular remodeling in both deficient proteins than in canonical G6PD (**Figure S5D**), independently supporting their increased conformational lability. Thus, A− combines near-normal baseline activity with accelerated functional decay, whereas Mediterranean G6PD combines impaired baseline activity with reduced thermal stability.

We next determined NADP⁺-bound cryo-EM structures of the canonical tetramer at 2.85 Å, A^−^ tetramer at 2.88 Å, Mediterranean tetramer at 2.94 Å, and Mediterranean dimer at 3.02 Å (**Figure 5F; Figures S6 and S7**), providing a high-resolution framework for interpreting these distinct biochemical phenotypes and a common reference for systematic comparison with published G6PD structures (**Figure S5D**). Of note, the novel NADP⁺-bound WT tetramer differed from the published unliganded structure by an all-atom RMSD of 3.75 Å, underscoring the value of a ligand-matched WT reference for distinguishing variant-associated structural changes from differences accompanying ligand occupancy (**Figure S5E**).

### Conformational heterogeneity at F188 destabilizes the Mediterranean tetramer

Unlike the well-resolved canonical and A^−^ reconstructions, the Mediterranean tetramer showed substantial conformational heterogeneity and reduced local density near the peripheral S188F substitution (**Figure 6A**). Three-dimensional variability analysis (3DVA)^51^ localized coordinated motion to the same region, identifying F188 as a focal point of tetramer instability (**Figure S5F**). To distinguish a protomer-intrinsic defect from oligomerization dependence, we also resolved the Mediterranean dimer at 3.02 Å. Its NADP⁺-binding region was well resolved and more homogeneous than in the tetramer, indicating that the S188F defect depends strongly on tetramerization (**Figure 6B; Figure S7**).

**Figure 6.**
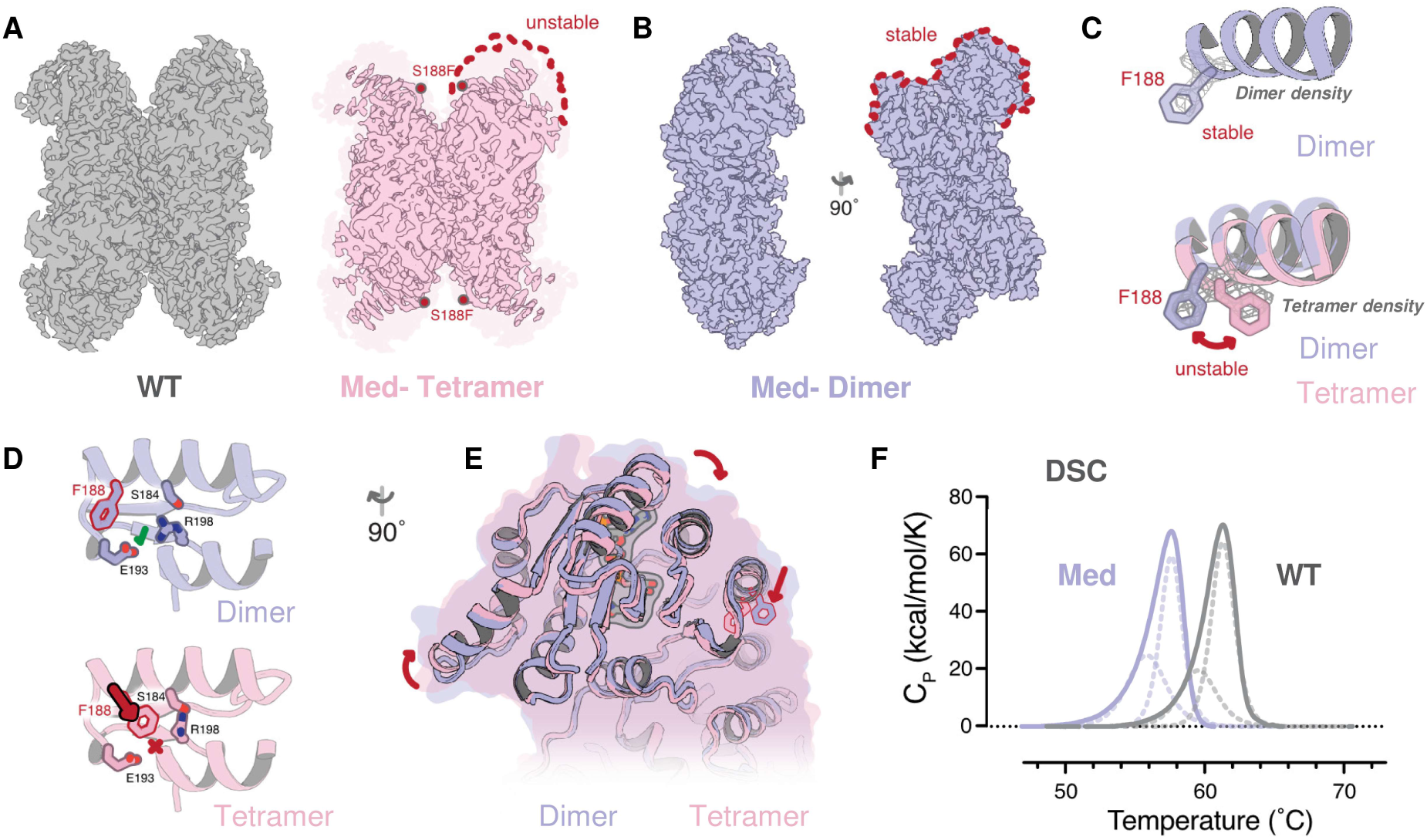
Oligomerization-dependent conformational heterogeneity destabilizes Mediterranean G6PD. Cryo-EM maps compare canonical and Mediterranean G6PD tetramers and localize reduced density around F188 in the variant **(A)**. The Mediterranean dimer is comparatively well resolved in this region **(B)**. Density comparisons show predominantly outward-facing F188 in the dimer and alternative conformations in the tetramer **(C)**. The associated S184–E193–R198 interaction network is remodeled **(D)**. Dimer/tetramer superposition reveals broader domain displacement and rotation **(E)**. Differential scanning calorimetry shows downward-shifted thermal transitions for Mediterranean G6PD **(F)**. See also Figures S5–S7.

F188 was predominantly outward-facing in the dimer, whereas the tetramer was consistent with alternative outward- and inward-facing conformations (**Figure 6C**). In the outward state, the local network, including the R198–E193 interaction, remained comparatively intact. Inward rotation instead displaced R198 from the protein core, moved the adjacent helix inward, and redirected R198 toward S184 (**Figure 6D**). Because S184 is itself altered in naturally occurring G6PD-deficiency variants, this region appears particularly functionally vulnerable. Thus, S188F couples alternative F188 conformations to remodeling of the surrounding S184–R198–E193 network rather than perturbing only its immediate residue environment.

This rearrangement propagated into displacement and rotation of the surrounding domain, linking F188 heterogeneity to broader oligomerization-dependent remodeling (**Figure 6E**). DSC provided orthogonal evidence of reduced thermal stability: relative to canonical G6PD, both Mediterranean transitions shifted downward by approximately 3.7°C, from 59.69 to 55.98°C and from 61.45 to 57.75°C (**Figure 6F**). These results agree with the fluorescence-based thermal transitions and distinguish Mediterranean G6PD from A^−^, whose transition temperatures remain near canonical despite pronounced time-dependent functional decay.

### A^−^ couples active-site remodeling to weakening of the dimer interface

The prominent difference between the A^−^ and WT structures was an elongation of the G6PD dimer. A^−^ extended approximately 4 Å farther at each end, increasing solvent-accessible surface area and reducing the buried protomer interface to ∼75% of WT (**Figures 7A–B**). Interprotomer contacts were also remodeled (**Figures 7C–D**), including weakening of D228–R219 and redistribution of the adjacent K275/E287/K290/D283 network (**Figure S5G**).

**Figure 7.**
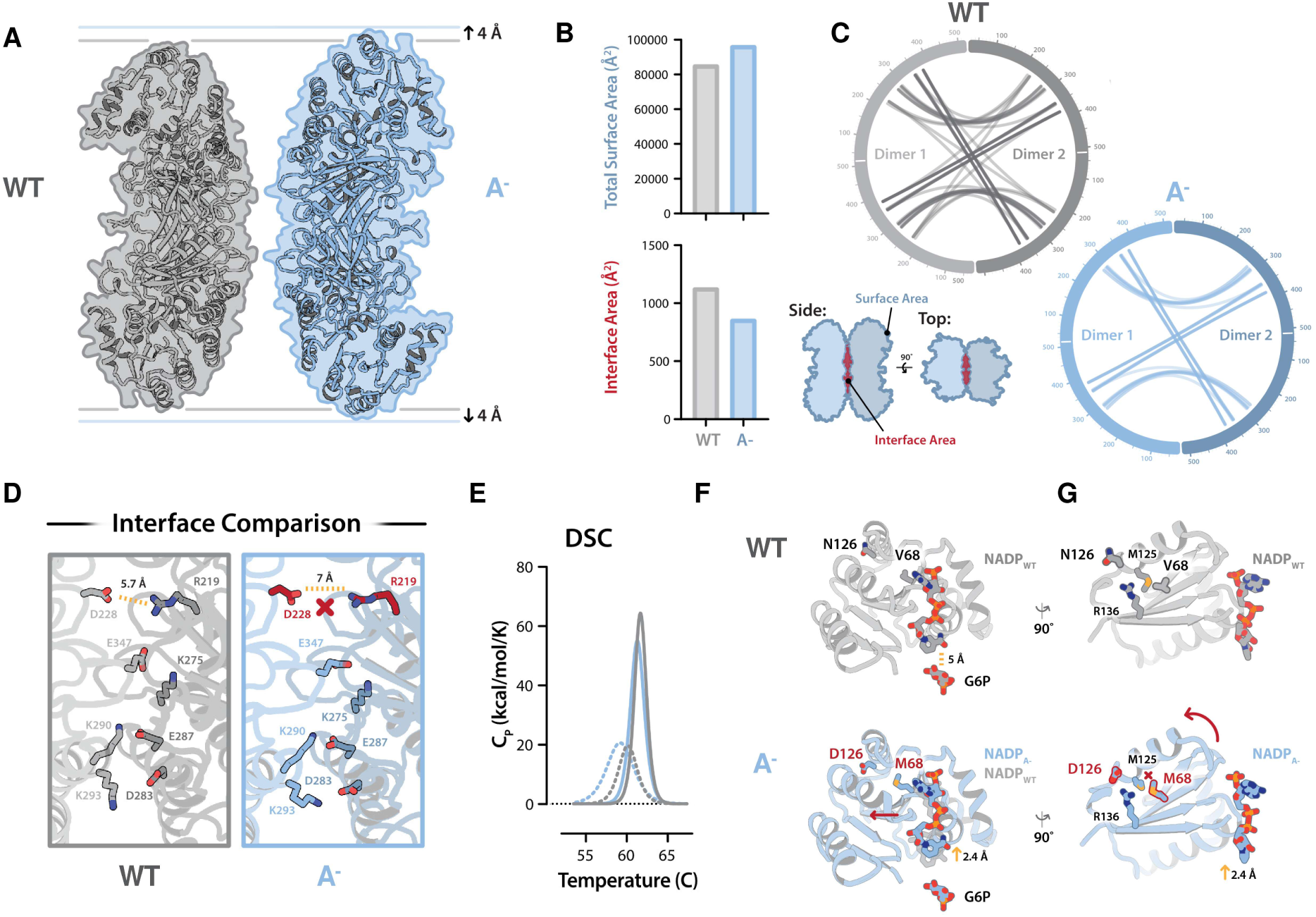
**The African A− variant remodels the G6PD dimer interface and catalytic NADP**⁺**-binding site.** NADP⁺-bound A− dimers are elongated by approximately 4 Å at each end relative to canonical G6PD **(A)**. Total solvent-accessible and buried interface areas are compared **(B)**. Interprotomer contact networks and representative interface interactions reveal reduced and remodeled intersubunit contacts **(C–D)**. Differential scanning calorimetry compares thermal transitions **(E)**. V68M remodels the catalytic region and displaces NADP⁺ relative to glucose-6-phosphate **(F)**. N126D alters the local interaction network and NADP⁺-contacting elements, further displacing catalytic NADP⁺ **(G)**. See also Figures S5 and S6.

DSC provided an orthogonal thermal-transition profile of this altered oligomeric state. Unlike Mediterranean G6PD, A^−^ showed only modest shifts in its first and second transitions relative to WT (58.60 versus 59.69°C and 61.01 versus 61.45°C). However, its first transition had lower apparent van’t Hoff enthalpy (172 versus 252 kcal mol^-1^) despite higher calorimetric enthalpy (104 versus 68 kcal mol^-1^), revealing altered energetic cooperativity not captured by transition temperature alone (**Figure 7E**). Thus, A^−^ weakens the dimer interface and alters transition cooperativity despite retaining near-normal baseline activity.

Interface remodeling accompanied structural changes around the catalytic NADP⁺-binding site. The bulkier V68M side chain imposed a steric constraint that displaced and rotated the neighboring helix (**Figure 7F**), while N126D formed a new polar interaction with R136 that stabilized the altered local conformation. These changes propagated to two NADP⁺-contacting loops: residues 109–115 shifted upward, and residues 140–146 moved laterally by approximately 2 Å (**Figure 7G**). Bound NADP⁺ was consequently displaced by ∼2.4 Å relative to WT and shifted away from the adjacent G6P-binding region. The resulting increase in hydride donor–acceptor distance could disfavor transfer from G6P to NADP⁺, providing a structural basis for progressive A− activity loss.^32,52^

Functional instability therefore converges across deficient alleles through distinct mechanisms: A^−^ retains greater baseline activity but develops kinetic fragility, whereas Mediterranean G6PD begins with lower activity, has lower thermal-transition temperatures, and accumulates dimers and trimers. These allele-specific defects connect *in vivo* enzyme decay to impaired RBC storage quality and transfusion efficacy.

## DISCUSSION

By resolving G6PD deficiency at the level of individual alleles, this study links inherited donor variation to stored-RBC biology and transfusion performance. African A^−^ (V68M/N126D) was associated with lower G6PD abundance, altered redox metabolism, greater oxidative and reduced osmotic hemolysis, reduced murine PTR, and smaller hemoglobin increments in humans. Mediterranean S188F instead combined low baseline activity and temperature-sensitive instability with greater post-transfusion plasma hemolysis and renal oxygenation deficits in mice. Allele-specific trajectories reconcile findings that appeared discordant when G6PD deficiency was treated as a single exposure.

The donor data place G6PD at a rate-limiting point in the storage response. Its abundance tracked with modeled PPP flux, whereas downstream PGD flux was comparatively independent of PGD abundance and appeared substrate constrained. The recurrent association of V68M/N126D with G6PD abundance, redox metabolites, and stress-induced hemolysis supports a progression from allele to protein dosage, constrained reducing capacity, and cellular injury. N126D alone, which defines A⁺ and is not deficient under the current WHO scheme,^2^ also associated with lower G6PD abundance and modest storage phenotypes. This does not recast A⁺ as deficient, but shows that an activity-based clinical category need not imply neutrality under prolonged *ex vivo* stress.

Two ancillary observations require bounded interpretation. Storage generated an N126D-like peptide signal in donors encoding Asn126. Deamidation can yield Asp and isoAsp,^53,54^ and the Asp-containing peptide is isobaric with genetically encoded N126D. While we neither isolated this proteoform nor established structural or functional equivalence to germline N126D, within the complete A^−^ structure we observed that D126 formed a new polar interaction with R136 consistent with stabilization of the altered local conformation. The present data do not isolate the function of the storage-derived Asp/isoAsp species or the independent contribution of N126D from V68M, but identify residue 126 as a genetic and post-translational node at which G6PD resilience may be modified.

Of note, in the REDS cohort, prevalence of the A-variant in donors of African descent overlaps with that of a ferroptosis-resistance allele set (STEAP3^10^, LPCAT3^10^, ACSL4^10^, FADS1/2^10^, GPX4^55^, p53^10^, SLC22A16^34^), contributing to a polygenic risk that predisposes mature RBC to oxidative hemolysis while conferring relative protection from osmotic fragility^56^. We hypothesize that this ancestry-linked constellation reflects balancing selection in malarial, hot, and dehydrating environments: traits that stiffen or dehydrate RBCs and remodel lipids may restrict osmotic lysis and parasite fitness, whereas co-segregating redox traits increase oxidative liability. Storage may expose the cost of this evolutionary trade-off.

The structures explain why variant identity matters. Early biochemical studies established interconverting G6PD oligomers and their stabilization by NADP⁺ and NADPH.^49,57^ Crystallography then defined the human fold and structural NADP⁺-binding site using the Canton variant and N-terminally truncated proteins,^5,58^ and structures of severe class I variants revealed long-range coupling among this site, oligomer interfaces, and the substrate-binding region.^6^ Cryo-EM later resolved sequence-intact WT G6PD without ligand, whereas ligand-bound tetramers used catalytically impaired D200N.^32^ The four near-atomic-resolution structures reported here therefore provide, to our knowledge, the first sequence-intact NADP⁺-bound WT tetramer and the first experimental structures of complete African A^−^ and Mediterranean G6PD. Matched ligand conditions distinguish variant- and oligomer-dependent remodeling from ligand-occupancy effects.

The structures reveal two failure modes. Molecular-dynamics simulations by Batista da Rocha et al.^59^ predicted that V68M and N126D together perturb the catalytic NADP⁺-binding domain more than either substitution alone; the A^−^ structure now resolves this cooperativity experimentally. Local remodeling propagates through NADP⁺-contacting loops, displaces catalytic NADP⁺, and elongates the dimer while reducing its interface, connecting near-normal initial activity to accelerated functional decay and showing why V68M alone is an incomplete proxy for A^−^.

Mediterranean S188F instead produces oligomerization-dependent heterogeneity: the ordered dimer favors an outward F188 conformation, whereas the tetramer samples states that remodel the S184–E193–R198 network and broader domain motion. This accords with lower baseline activity, reduced apparent thermal stability in irreversible oligomerization-coupled transitions, and NADP⁺-dependent stabilization, providing a structural counterpart to preserved refrigeration storage followed by stress-induced hemolysis after rewarming^26^. Oligomer locking can improve variant function; dimer locking activated A^−^ whereas tetramer locking activated Mediterranean G6PD.^7,60^ These alleles may therefore require different stabilization strategies.

The mouse studies connect these defects to RBC aging and recipient physiology. Biotin labeling showed age-dependent loss of G6PD abundance and activity, with variant-specific kinetics consistent with classic measurements of shorter enzyme half-lives in A^−^ and Mediterranean RBCs.^48^ A^−^ RBCs developed greater oxidative dysfunction and lower PTR, whereas Mediterranean RBCs retained comparatively preserved PTR, but produced stronger plasma hemolysis signatures and the largest renal oxygenation deficit after controlled hemorrhage and serial isovolemic exchange, highlighting a disconnect between Mediterranean RBC circulatory capacity and preserved function that neither PTR or HbA/HbS tracing can capture. This supports a circulating-cell-injury-dominant A− phenotype and a more hemolysis/dysfunction-dominant Mediterranean phenotype, although PTR does not distinguish hemolysis, extravascular clearance, and transient sequestration.

Human data extend this framework across complementary clinical settings. In 4,464 single-unit transfusion events, African A− was associated with allele-dose-dependent reductions in adjusted hemoglobin increment; Mediterranean units were too uncommon for comparable estimates. Francis et al. directly measured autologous ^51^Cr recovery in healthy donors,^17^ whereas Sagiv et al. inferred allogeneic persistence in SCD from bulk HbA across serial, multi-unit transfusions.^19^ Here, a single ^51^Cr-labeled allogeneic study unit was followed from minutes through four weeks in recipients with SCD while all other exchange units were G6PD sufficient. Lower recovery and survival were accompanied by hemolysis and sustained multi-omic remodeling, agreeing with the larger recipient-linked analysis and chromium-free murine PTR. Its value is mechanistic specificity rather than a precise population-effect estimate.

The magnitude is modest per unit but potentially biologically meaningful. A packed RBC unit contains approximately 2.0–2.5×10^12^ cells; the observed 8.12- at 24h and 11.75-percentage-point differences at four weeks correspond to approximately 1.6–2.0×10^11^ and 2.4–3.0×10^11^ fewer donor RBCs represented in circulation, respectively - ∼0.8–1.4 days of normal whole-body RBC turnover in an adult. Assuming an expected increment of ∼1 g/dL per unit, these differences represent ∼0.08–0.12 g/dL less effective hemoglobin increment per affected unit, or 0.16–0.36 g/dL if a comparable effect occurred across two or three deficient units. At ∼55 g hemoglobin per unit, they also represent ∼6–9 mL less maximal oxygen-carrying capacity in circulation per affected unit, consistent in direction with the renal oxygenation deficit in mice. Separately, if the measured hemolysis difference reflects intravascular release, it corresponds to ∼0.02–0.04 g heme and 1–4 mg iron per unit. While PTR itself does not establish the route of RBC loss, elegant work by Roussel et al.^61^ previously linked PTR to splenic sequestration and extravascular – rather than intravascular - hemolysis. On the other hand, no differences in end-of-storage spontaneous hemolysis were here observed either in mice or REDS donors, where *G6PD* alleles only tracked with higher oxidative hemolysis and lower osmotic fragility, with no association to storage hemolysis, consistent with prior studies.^24,26^ These physiological-scale translations are not clinical endpoints, but recurrent shortfalls could increase ineffective transfusion and exposure to hemolysis products, contributing to additional-unit requirements, donor exposure, iron loading, and alloimmunization risk.^62,63^

The donor exposome - the aggregate of dietary, occupational, recreational, and medical exposures - may broaden this principle beyond inherited deficiency.^64^ Caffeine, consumed daily by more than 67% of adults,^65^ is especially relevant: 50 μM caffeine, a concentration attainable after approximately 1–2 cups of coffee, acutely reduced recombinant canonical G6PD activity by approximately 33%^66^ but did not further reduce African A− activity.^67^ Environmental exposures may therefore constrain G6PD-dependent redox buffering in genetically sufficient donors while interacting differently with individual alleles. However, caffeine is unlikely to explain the SCD pilot trial PTR results. Intake was not prospectively recorded, but neither caffeine (5h half-life in circulation) nor paraxanthine was detected in transfused units or recipient plasma or RBCs sampled 2h after transfusion. Exposure outside the sampling window or below detection cannot be excluded, but the available measurements and preferential inhibition of canonical but not A-G6PD^67^ make caffeine more likely to attenuate than generate the deficient-versus-sufficient contrast.

Several limitations define the scope of inference. The vein-to-vein analysis is observational, uncommon alleles - particularly Mediterranean S188F - were insufficiently represented for human outcome estimates, and X-linked mosaicism complicates inference in heterozygous females. The prospective SCD cohort was exploratory, but directly measured one allogeneic unit and converged with the larger human analysis and chromium-free murine experiments. A genotype-by-^51^Cr interaction cannot be completely excluded, although it is unlikely to explain that cross-platform agreement. The humanized alleles share a murine promoter and regulatory context, and the sensitized storage and exchange paradigms model selected aspects of human transfusion rather than reproducing them quantitatively. These constraints limit extrapolation and effect-size precision, but not the internally concordant conclusion that variant-resolved G6PD status influences stored-RBC biology and post-transfusion performance.

The evidence supports a practical middle ground: G6PD-deficient units are not uniformly unsafe, and remain fully acceptable for most indications, but neither are they biologically equivalent to G6PD sufficient units under all storage and recipient conditions. Proportionate mitigation could include recording known A^−^ or Mediterranean status, prioritizing G6PD-sufficient units when durable efficacy or tolerance of hemolysis is especially important - such as chronic or exchange transfusion in SCD, neonatal transfusion, and massive transfusion or hemorrhage - and evaluating program-level or one-time, variant-resolved screening at first donation. Such testing is not operationally unprecedented: some blood systems already screen for or restrict donation according to G6PD status,^23^ and expanding genomic platforms for extended blood-group typing^68^ could incorporate donor-quality alleles^69^ without a separate assay for every donation. Implementation should be variant resolved and not ancestry based, with focused pilots defining effects on supply, workflow, cost, and clinical outcomes before scale-up. Within precision transfusion medicine,^70^ this is a fitness-for-purpose strategy rather than a judgment that deficient blood is categorically unusable, which would pose unreasonable and unjustified constraints on the inventories of blood banks serving regions where G6PD variant is prevalent (e.g., 37.5% in Sokoto, Nigeria).^71^

Beyond its implications for transfusion practice, the allele-resolved structural and functional framework established here should prove broadly useful to biologists and clinicians investigating dysregulated pentose phosphate pathway biology - an increasingly recognized axis of redox control in aging, immunity, neurodegeneration, obesity, cardiovascular disease, and cancer.^3^

## Online METHODS

### Recipient Epidemiology and Donor evaluation Study (REDS) RBC Omics

A detailed description of the study design, enrollment criteria and main outcomes has been provided in previous publications,^27,72^ and – while redundant with the literature - will be included below with significant overlap with published methods.

### Donor recruitment in the REDS RBC Omics study

#### Index donors

A total of 13,758 donors were enrolled in the Recipient Epidemiology and Donor evaluation Study (REDS) RBC Omics at four different blood centers across the United States (https://biolincc.nhlbi.nih.gov/studies/reds_iii/). Of these, 97% (13,403) provided informed consent and 13,091 were available for metabolomics analyses in this study – referred to as “index donors”. A subset of these donors were evaluable for hemolysis parameters, including spontaneous (n=12,753) and stress (oxidative and osmotic) hemolysis analysis (n=10,476 and 12,799, respectively) in ∼42-day stored leukocyte-filtered packed RBCs derived from whole blood donations from this cohort ^73^. Methods for the determination of FDA-standard spontaneous (storage) hemolysis test, osmotic hemolysis (pink test) and oxidative hemolysis upon challenge with AAPH have been extensively described elsewhere ^56^.

#### Recalled donors

A total of 643 donors scoring in the 5^th^ and 95^th^ percentile for hemolysis parameters at the index phase of the study were invited to donate a second unit of pRBCs, a cohort henceforth referred to as “recalled donors”. These units were assayed at storage days 10, 23 and 42 for hemolytic parameters and mass spectrometry-based high-throughput metabolomics ^74^, proteomics ^75^, lipidomics ^76^ and ICP-MS analyses ^77^. Under the aegis of the REDS-IV-P project ^27^, a total of 1,929 samples (n=643, storage day 10, 23 and 42) were processed with this multi-omics workflow. Of the 643 parent units, 638 were genotyped for 362 G6PD SNPs.

#### mQTL and pQTL studies

The workflow for the mQTL analysis of metabolites and pQTL analysis for G6PD protein levels in REDS Index and Recalled donors is consistent with previously described methods from our pilot mQTL study on 250 recalled donors^78^ and follow up studies on specific metabolic pathways (e.g., glycolysis^79^) other proteins (e.g., GPX4^55^) on the full index cohort. Details of the genotyping and imputation of the RBC Omics study participants have been previously described by Page, et al.^24^ Briefly, genotyping was performed using a Transfusion Medicine microarray^69^ consisting of 879,000 single nucleotide polymorphisms (SNPs); the data are available in dbGAP accession number phs001955.v1.p1. Imputation was performed using 811,782 SNPs that passed quality control. After phasing using Shape-IT ^80^, imputation was performed using Impute2 ^81^ with the 1000 Genomes Project phase 3 ^81^ all-ancestry reference haplotypes. We used the R package SNPRelate ^82^ to calculate principal components (PCs) of ancestry. We performed association analyses using an additive SNP model in the R package ProbABEL^83^ and 643 REDS Recalled donor study participants who had G6PD protein levels and imputation data on serial samples from stored RBC components that passed respective quality control procedures. We adjusted for sex, age (continuous), frequency of blood donation in the last two years (continuous), blood donor center, and ten ancestry PCs. Statistical significance was determined using a p-value threshold of 5x10^-8^. We only considered variants with a minimum minor allele frequency of 1% and a minimum imputation quality score of 0.80. The OASIS: Omics Analysis, Search & Information a TOPMED funded resources ^84^, was used to annotate the top SNPs. OASIS annotation includes information on position, chromosome, allele frequencies, closest gene, type of variant, position relative to closest gene model, if predicted to functionally consequential, tissues specific gene expression, and other information.

The 105 compound heterozygotes, representing 0.82% of the genotyped population, which were retained in all variant-level analyses. SNP association tests were performed using an additive allele dosage model at each tested SNP rather than categorical genotype groups. Estimates should therefore be interpreted as marginal, SNP-specific associations rather than independent effects of compound genotypes. G6PD SNP genotype dosages were interpreted according to their X-linked inheritance, treating a hemizygous male as equivalent to a homozygous female (i.e., dosages of 0 or 2), within the association pipeline described above. Separate analyses showed that allele A^+^ association to lower protein levels and protection from osmotic fragility persisted after excluding donors carrying V68M in cis.

#### Determination of hemoglobin and bilirubin increment via the vein-to-vein database

Association of G6PD SNPs with hemoglobin increments was performed by interrogating the vein-to-vein database, as described in Roubinian et al.^25^. Analyses were performed on the whole REDS population and further stratified for caffeine measurements, with a focus on the upper 50^th^ percentile of caffeine measurements in REDS donors.

#### Vein-to-vein database: General Study Design

We conducted a retrospective cohort study using electronic health records from the National Heart Lung and Blood Institute (NHLBI) Recipient Epidemiology and Donor Evaluation Study-III (REDS-III) program available as public use data through BioLINCC ^85,86^. The database includes blood donor, component manufacturing, and patient data collected at 12 academic and community hospitals from four geographically diverse regions in the United States (Connecticut, Pennsylvania, Wisconsin, and California) for the 4-year period from January 1, 2013 to December 31, 2016. Genotype and metabolomic data from the subset of blood donors who participated in the REDS-III RBC-Omics study ^87^ was linked to the dataset using unique donor identifiers.

#### Study Population and Definitions

Available donor genetic polymorphisms for G6PD were linked to issued RBC units using random donor identification numbers. Among transfusion recipients, we included all adult patients who received a single RBC unit during one or more transfusion episodes between January 1, 2013 and December 30, 2016. Recipient details included pRBC storage age, and blood product issue date and time. We collected hemoglobin levels measured by the corresponding clinical laboratory prior to and following each RBC transfusion event (0h and 24h after transfusion).

#### Transfusion Exposures and Outcome Measures

All single RBC unit transfusion episodes linked to genetic polymorphism for G6PD were included in this analysis. A RBC unit transfusion episode was defined as any single RBC transfusion from a single donor with both informative pre- and post-transfusion laboratory measures and without any other RBC units transfused in the following 24hour time period. The outcome measures of interest were change in hemoglobin (ΔHb; g/dL) following a single RBC unit transfusion episode. These outcomes were defined as the difference between the post-transfusion and pre-transfusion levels. Pre-transfusion thresholds for these measures were chosen to limit patient confounding (e.g., underlying hepatic disease). For pre-transfusion hemoglobin, the value used was the most proximal hemoglobin measurements prior to RBC transfusion, but at most 24 hours prior to transfusion. Furthermore, we excluded transfusion episodes where the pre-transfusion hemoglobin was greater than 9.5 g/dL, and the hemoglobin increment may be confounded by hemorrhage events. For post-transfusion hemoglobin, the laboratory measure nearest to 24hours post-transfusion, but between 12- and 36-hours following transfusion was used.

### Experimental mouse models

#### Humanized G6PD deficient mice

Human genomic G6PD sequence spanning exons 3–13 was substituted into the endogenous murine G6pd locus, whereas murine exon 2 was retained to avoid perturbing the overlapping IKKγ coding region.^88–90^ The endogenous murine promoter and non-targeted regulatory context were also retained, while avoiding retention of a 13 bp scar in the region between the murine promoter and the starting codon that was present in previous humanized G6PD deficient canonical and Med-strains^28^; thus, the models humanize the G6PD genomic/coding interval but not the promoter or enhancer landscape. All three lines were studied on the same C57BL/6J background, allowing protein-abundance differences to be interpreted as allele-dependent effects on expression and/or protein stability within a shared murine regulatory environment. Wild-type C57BL/6J mice were included as controls and omitted from selected panels only when their results overlapped those of the humanized canonical line.

#### Storage and Post-transfusion recovery (PTR) studies in humanized G6PD deficient mice

Mouse post-transfusion recovery (PTR) studies were performed as previously described ^91^. Humanized mice expressing canonical G6PD or the deficient variants A- (V68M; N126D) and Med-(S188F) were previously described in physiology studies^88–90^, but never before tested in the context of RBC storage. Storage of humanized G6PD canonical, A- and Med-deficient mice (n=3) for 0, 1, 3, 7 and 14 days (Experiment 1) or 0 and 12 days (Experiment 2). End of storage of day 12 samples was followed by transfusion into Ubi-GFP mice, which were used as recipients to allow visualization of the test cells in the non-fluorescent gate. To control for differences in transfusion and phlebotomy, mCherry (red fluorescent-labeled) RBCs were used as a tracer RBC population (never stored). These RBCs were added to stored RBCs immediately prior to transfusion. PTR was calculated by dividing the post-transfusion by the pre-transfusion ratio (Test/Tracer), with the maximum value set as 1 (or 100% PTR).

#### In vivo biotinylation studies

In vivo biotinylation studies of murine RBCs, followed by flow-sorting of biotin+ and biotin-cells was performed as described by Hoffmann-Fezer et al.^92^, as modified by Zimring et al.^93^. Flow-sorted RBCs underwent standard G6PD activity assay and mass spectrometry-based metabolomics was performed as previously described.^88^

### Mouse transfusion and tissue oxygenation

#### Donor mouse blood collection and storage

G6PD sufficient and deficient mice were anesthetized (induction 3-4.5% isoflurane and 1.5-3% in 100% oxygen for maintenance). A V-shaped cut was made through the skin and abdominal wall about 1 cm caudal to the last rib using a sterile micro dissecting scissors and forceps. A 23 Guage needle 1” attached to a 1 ml syringe was inserted through the diaphragm and into the vena cava to exsanguinate the animal. Blood from donor mice was pooled and processed into red blood cells and stored in Citrate-Phosphate-Dextrose-Adenine solution for 7 days under refrigeration (4-6C^○^).

#### Surgical procedures

Recipient C57BL6J mice (n=5/group) were anesthetized (induction 3-4.5% isoflurane and 1.5-3% for maintenance in 21% oxygen). Vital signs including heart rate, respiratory rate, pulse oximetery and rectal temperature were monitored using a PhysioSuite MouseSTAT and RightTemp Module-based system (Kent Scientific Corporation) continuously and recorded every 15 min. carotid artery cannulation: A transverse incision (∼1 cm) over the trachea. Using a hemostat, the omohyoid muscle was bluntly dissected longitudinally to expose the left carotid artery. A 3mm section of the vessel was isolated and a small incision was made. A mouse carotid arterial catheter (https://www.instechlabs.com /products/catheters/mouse-carotid) was inserted towards the heart with the assistance of the micro dissecting hook and forceps. A smooth needle holder without lock was used to hold the portion of the catheter inside the vessel tightly before removing the micro bulldog clamp. The catheter was advanced with a pair of forceps while loosening the needle holder slowly until the silicone collar segment of catheter was placed in the vessel. The loose caudal ligature was tied around the catheter and vessel to secure, but not to occlude, the catheter. The catheter was filled with heparin lock solution (10 U/mL of heparin) to maintain patency and used to perform blood removal and replacement.

#### Sequential exchange transfusion

C57BL6J mice (25-30g) underwent a sequential exchange transfusion by removing ∼200 μL of blood through the carotid artery catheter and replacing with an equal volume of either (1) humanized canonical nondeficient G6PD (HuCan), or African (A-) or Mediterranean (Med-) strains. 7-day refrigerator stored mouse donor blood prepared at 45% hematocrit in 5% human serum albumin. Blood withdrawal and transfusions were performed manually over a 5-minute period for each exchange followed by a 10-minute pO2 measurement interval. Note that this process is done in a stepwise fashion with blood removal followed immediately by RBC/albumin administration. This process was repeated until the target number 7 exchanges were performed.

#### Imaging

Prior to imaging, 71 mmol (∼0.15 mL) of OX071was administered through the carotid catheter. The signal amplitude and pO2 maps were obtained with IRESE acquisition protocol with the following parameters: π/2 and π pulse lengths of 60 ns, 16-step phase-cycling scheme, spin-echo delay of 400 ns, equal solid angle spaced 208 projections, 22 baselines, 0.75 G/cm maximum gradients, 10 logarithmically spaced time delays (410 ns to 40 μs), 55 μs TR, and image-acquisition time of 2 min. Three images were collected for over 10 minutes for each experimental phase (before exchange transfusion, middle of exchange transfusion (after 4, exchange volumes ∼ 0.2 ml), end of exchange transfusion (after 7 total exchange volumes ∼2 ml). IRESE images were reconstructed using filtered back projection (FOV = 4.24 cm) in isotropic 64 × 64 × 64 voxels with 0.66 mm × 0.66 mm × 0.66 mm voxel size. The R1 (1/T1) values were calculated using a single exponential fit after selecting the highest intensity voxels with a thresholding filter (set at 3% of the maximum amplitude). R1 maps were converted to pO2 using a predetermined calibration (slope: 109.67 torr/Ms−1, intercept: 0.124 Ms−1).

### Image analysis

Kidney regions of interest (ROIs) were generated automatically using a three-dimensional U-Net (UNet3D) deep learning model implemented in PyTorch. The network was optimized with a Dice–cross entropy composite loss until convergence to generate the final model weights. The model weights are then utilized by a custom python script that normalizes the EPRI amplitude image input and passes the image through the network to produce a probability map of the kidney tissue. This probability map is threshholded (0.5) and the two largest connected components are extracted as candidate kidneys and saved as a binary mask. The binary kidney mask is delineated by comparing their centroids along the left–right axis to assign consistent anatomical labels. Each mask is stored as a ROI-slave image in a project file compatible with the MATLAB-based GUI, ArbuzGUI.

Data is presented as the median and minimum to maximum pO2 mmHg for all images (∼15 per group) obtained at each time point. An ANOVA with was performed a multiple comparisons test in GraphPad Prism 10.6.1 (Boston, MA, UA). P Values are noted for each comparison with statistical significance set at P < 0.05 at a 95% CI.

### Clinical trial

#### Transfusion of G6PD deficient RBCs in patients with SCD

We performed a single blind clinical study in adults with SCD who were receiving monthly chronic automated exchange transfusions (The Impact of Oxidative Stress on Erythrocyte Biology (RBC Survival) ClinicalTrials.gov ID: NCT04028700).

NCT04028700 was designed as a participant-blinded, two-treatment, two-period, two-sequence crossover study targeting 16 participants who would complete both periods. Treatment sequence was determined by the availability of a compatible qualifying unit rather than by formal randomization. Recruitment was interrupted and the study was terminated during the COVID-19 pandemic before the planned enrollment and crossover were completed. The final cohort comprised eight evaluable participants, each contributing one study transfusion: five received a G6PD-sufficient study unit and three a G6PD-deficient study unit. The completed study therefore did not retain the paired design or precision assumed in the prespecified power calculation and was analyzed as an exploratory between-group comparison.

All units for the exchange were tested for G6PD activity, and all units were from sickle trait-negative, ABO-compatible, CEK-matched, and cross-match compatible donors. All units were stored in AS-1, had comparable donor hemoglobin (21.4 ± 1.9 g/dL) and storage age at transfusion (22.8 ± 6.3). Subjects were blinded to the G6PD infusion assignment (G6PD normal vs. G6PD deficient red cells). All units used in the exchange were G6PD-normal except for one study RBC unit, the final unit transfused. During the last 30-45 minutes (+/- 30 minutes) of the exchange, a 50mL sterile sample was removed from the final unit of the exchange (the experimental RBC unit); one aliquot was used for ^51^Cr-labeling and the other for metabolomic and lipidomic studies. Once the exchange was complete, the subject was then infused with the prepared ^51^Cr-labeled aliquot per established, Radioactive Drug Research Committee reviewed and approved nuclear medicine protocols. Blood samples from the patient were obtained pre-infusion, and at approximately 5, 7.5, 10, 12.5, 15, 30 minutes; 1, 24, 48, 72 hours; and 1, 2, 3, 4 weeks post-infusion. We measured the post-transfusion recovery (PTR) and 4 week survival of the labeled donor RBCs, accounting for the expected elution of ^51^Cr using standard formulas. A CBC, hemoglobin-thalassemia profile, bilirubin (total and direct), Ferritin, C-reactive protein, and lactate dehydrogenase were measured pre-transfusion and at 1 hour, 1 day, and 1,2,3, and 4 weeks post-infusion to explore their correlation with the ^51^Cr results.

For 24h ^51^Cr post-transfusion recovery and four-week labeled-RBC survival, group means and deficient-minus-sufficient mean differences were reported with two-sided 95% Welch confidence intervals. Exact one-sided Mann–Whitney tests evaluated the prespecified directional hypothesis of lower recovery and survival for G6PD-deficient units. Given the incomplete crossover and small final cohort, nominal P values were treated as exploratory descriptors; the confidence intervals provide the primary statement of precision. Accordingly, we emphasize the observed direction and magnitude of the effects and retain the one-sided P values only as exploratory descriptors, interpreting these findings as consistent with, rather than definitive estimates of, the signals observed independently in the larger linked donor–recipient cohort and murine studies.

G6PD-sufficient and G6PD-deficient units underwent identical standard ^51^Cr-labeling and sampling procedures. Although a genotype-by-label interaction cannot be formally excluded, the directionally concordant findings from the linked donor–recipient analysis and fluorescence-based murine PTR experiments - neither of which involved chromium - argue against a labeling-specific effect as the sole explanation. These convergent findings nevertheless require validation in a larger, prospectively allocated clinical study. The implications of this limitation are considered in the Discussion.

##### Omics analyses

Omics methods described below follow exactly those described in previous publications, as referenced in each section that follows.

##### High-throughput metabolomics

Metabolomics extraction and analyses in 96 well-plate format were performed as described, with identical protocols for human or murine RBCs^94,95^. RBC samples were transferred on ice on 96 well plate and frozen at -80 °C at Vitalant San Francisco (human RBCs) or University of Virginia (murine RBCs) prior to shipment in dry ice to the University of Colorado Anschutz Medical Campus. Plates were thawed on ice then a 10 uL aliquot was transferred with a multi-channel pipettor to 96-well extraction plates. A volume of 90 uL of ice cold 5:3:2 MeOH:MeCN:water (v/v/v) was added to each well, with an electronically-assisted cycle of sample mixing repeated three times. Extracts were transferred to 0.2 µm filter plates (Biotage) and insoluble material was removed under positive pressure using nitrogen applied via a 96-well plate manifold. Filtered extracts were transferred to an ultra-high-pressure liquid chromatography (UHPLC-MS — Vanquish) equipped with a plate charger. A blank containing a mix of standards detailed before ^96^ and a quality control sample (the same across all plates) were injected 2 or 5 times each per plate, respectively, and used to monitor instrument performance throughout the analysis. Metabolites were resolved on a Phenomenex Kinetex C18 column (2.1 x 30 mm, 1.7 um) at 45 °C using a 1-minute ballistic gradient method in positive and negative ion modes (separate runs) over the scan range 65-975 m/z exactly as previously described.^94^ The UHPLC was coupled online to a Q Exactive mass spectrometer (Thermo Fisher). The Q Exactive MS was operated in negative ion mode, scanning in Full MS mode (2 μscans) from 90 to 900 m/z at 70,000 resolution, with 4 kV spray voltage, 45 sheath gas, 15 auxiliary gas. Following data acquisition, .raw files were converted to .mzXML using RawConverter then metabolites assigned and peaks integrated using ElMaven (Elucidata) in conjunction with an in-house standard library^97^.

##### Lipid hydroperoxides and glutathionylated lipids: sample preparation

Oxylipins were extracted via a modified protein crash from previously described.^10,14,16^ Extraction of oxylipins from red blood cells were as follows: 10 µL of red cells was aliquoted directly into 90 μL of MeOH:MeCN:H_2_O (5:3:2, v:v:v) and pipetted up and down 5x per sample. Samples were then vortexed at 4 °C for 30 minutes. Following vortexing, samples were centrifuged at 12700 RPM for 10 minutes at 4 °C and 80 μL of supernatant was transferred to a new tube for analysis. 10 μL of extract from each sample was also combined to create a technical mixture, injected throughout the run for quality control. Lipidomics were extracted using identical prepartion techniques as oxylipins, save for utilizing MeOH:IPA (1:1, v:v) instead of MeOH:MeCN:H_2_O (5:3:2, v:v:v) as an extraction buffer.

##### High-throughput Oxylipin Analysis

Analyses were performed as previously published via a modified gradient optimized for the high-throughput analysis of oxylipins.^10,14,16^ Briefly, the analytical platform employs a Vanquish UHPLC system (Thermo Fisher Scientific, San Jose, CA, USA) coupled online to a Q Exactive mass spectrometer (Thermo Fisher Scientific, San Jose, CA, USA). Lipid extracts were resolved over an ACQUITY UPLC BEH C18 column (2.1 x 100 mm, 1.7 µm particle size (Waters, MA, USA) using mobile phase (A) of 20:80:0.02 ACN:H_2_O:FA and a mobile phase (B) 20:80:0.02 ACN:IPA:FA. For negative mode analysis the chromatographic the gradient was as follows: 0.35 mL/min flowrate for the entire run, 0% B at 0 min, 0% B at 0.5 min, 25%B at 1 min, 40%B at 2.5min, 55% B at 2.6min, 70% B at 4.5 min, 100% B at 4.6 min, 100% B at 6 min, 0% B at 6.1 min and 0% B at 7 min. The Q Exactive mass spectrometer (Thermo Fisher) was operated in negative ion mode, scanning in Full MS mode (2 μscans) from 150 to 1500 m/z at 70,000 resolution, with 4 kV spray voltage, 45 sheath gas, 15 auxiliary gas. Calibration was performed prior to analysis using the Pierce^TM^ Positive and Negative Ion Calibration Solutions (Thermo Fisher Scientific).

##### High-throughput Lipidomic Analysis

Lipid extracts were analyzed (10 µL per injection) on a Thermo Vanquish UHPLC/Q Exactive MS system using a previously described^76^ 5 min lipidomics gradient and a Kinetex C18 column (30 x 2.1 mm, 1.7 µm, Phenomenex) held at 50 °C. Mobile phase (A): 25:75 MeCN:H_2_O with 5 mM ammonium acetate; Mobile phase (B): 90:10 IPA:MeCN with 5 mM ammonium acetate. The gradient and flow rate were as follows: 0.3 mL/min of 10% B at 0 min, 0.3 mL/min of 95% B at 3 min, 0.3 mL/min of 95% B at 4.2 min, 0.45 mL/min 10% B at 4.3 min, 0.4 mL/min of 10% B at 4.9 min, and 0.3 mL/min of 10% B at 5 min. Samples were run in positive and negative ion modes (both ESI, separate runs) at 125 to 1500 m/z and 70,000 resolution, 4 kV spray voltage, 45 sheath gas, 25 auxiliary gas. The MS was run in data-dependent acquisition mode (ddMS^2^) with top10 fragmentation. Raw MS data files were searched using LipidSearch v 5.0 (Thermo).

##### Statistical analysis

Acquired data was converted from raw to mzXML file format using Mass Matrix (Cleveland, OH, USA). Analysis was done using MAVEN, an open-source software program for oxylipin analysis. Lipid assignments and peak integration were performed using LipidSearch v 5.0 (Thermo Fisher Scientific). Samples were analyzed in randomized order with a technical mixture injected interspersed throughout the run to qualify instrument performance. Data analysis and Statistical analyses – including hierarchical clustering analysis (HCA), linear discriminant analysis (LDA), uniform Manifold Approximation and Projection (uMAP), correlation analyses and Lasso regression were performed using both MetaboAnalyst 5.0 and RStudio (2024.12.1 Build 563).

#### Proteome-constrained modeling of human red blood cells

Proteomic data for RBCs obtained from 638 donors of the REDS Recall cohort and stored for 10, 23, and 42 days were collected as previously described^98^. Protein copy numbers were computed via the “total protein approach” ^99–101^ using the mean corpuscular hemoglobin values calculated from the CBC data obtained at the time of donation for the corresponding donor. Samples were scaled assuming hemoglobin comprised 95% of the proteome dry weight and mapped onto the human RBC genome-scale metabolic reconstruction (RBC-GEM version 1.2.0)^102^. A total of 1877 proteome-constrained models for sample were subsequently derived by following the OVERLAY workflow as previously described in detail ^102,103^. Individual protein constraints were relaxed by 3%, and the global relaxation constraint was minimized to its smallest possible value that allowed for solution feasibility in simulations.

#### Proteome-constrained modeling of mouse red blood cells

The human RBC-GEM (version 1.2.0)^102^ and the vertebrate homology data from Mouse Genome Informatics database (MGI version 6.24)^104^ were utilized to create a mouse-specific RBC-GEM by mapping human genes to their mouse orthologs. Proteomic data for RBCs from 12 humanized mouse models expressing G6PD canonical non-deficient protein, 12 mice expressing African V68M variant, and 12 mice expressing the Mediterranean S188F variant were collected as described above. Protein copy numbers were computed via the “total protein approach”^99–101^ using a mean corpuscular hemoglobin value of 13.9 picograms.^105^. Samples were scaled assuming hemoglobin comprised 95% of the proteome dry weight, and 36 proteome-constrained models were subsequently derived for each sample by following the OVERLAY workflow as previously described in detail,^102,103^. Individual protein constraints were relaxed by 3%, and the global relaxation constraint was minimized to its smallest possible value that allowed for solution feasibility in simulations.

#### Analysis of proteome-constrained models

Proteome-constrained models were simulated using flux balance/flux variability analysis (FBA/FVA) to determine maximum fluxes, maximum enzyme abundances, and effective flux ranges. Spearman rank correlation coefficients (ρ) were computed between maximum flux and enzyme abundance to classify reactions as based on their simulated protein abundance as abundance-dependent (ρ ≥ 0.8), abundance-correlated (0.5 ≤ ρ < 0.8), or abundance-independent (ρ < 0.5). Only reactions that carried flux were considered. Within each classification, reactions were grouped by metabolic category.

Statistically significant differences for simulated flux ranges were determined within the group of 638 human donors. For each donor, the mean flux range for each flux-carrying reaction were determined. Flux ranges were Z-standardized across donors, then divided into three groups based on number of donor allele copies for African V68M G6PD variant with hemizygous males considered part of the group with two copies. The Kruskal-Wallis H-test was then conducted to identify reactions with statistically significant (p < 0.0002) differences in flux range within the three groups.

Statistically significant differences for simulated flux ranges were also determined within the group of 36 mouse models. Flux ranges for each flux-carrying reaction were Z-standardized across the mouse models, then divided into three groups based on the expressed G6PD protein variant. The Kruskal-Wallis H-test was then conducted to identify reactions with statistically significant (p < 0.05) differences in flux ranges within the three groups.

All simulations performed were implemented in Python 3.11 using the COBRApy package (version 0.29.1)^106,107^ and its implementation of algorithms for FBA/FVA.^108^ Statistical analyses of simulation results were performed using the SciPy package (version 1.15.3) .^109^ Statistically significant differences in flux ranges between were visualized as a heatmap using the Morpheus online software tool (https://software.broadinstitute.org/morpheus, version 1.0.18 accessed 7/27/25) with the mean flux range values of each G6PD group displayed. Area plots of flux ranges and min-max normalized enzyme abundances across G6PD groups were visualized using Matplotlib (version 3.10.3).^110^

#### Recombinant G6PD protein expression and purification

Human G6PD, WT, Med-(S188F), and A-(V68M, N126D) were expressed from a pET28 vector with an N-terminal 6xHis-TEV-Thrombin sequence (GenScript). Transformed C41 cells (LGC Biosearch Technologies) were grown at 37 °C overnight in LB from a single colony. The starter culture was used to inoculate TB at a 1:25 ratio. The cultures were grown at 37 °C and 220 rpm until an OD_600_ of 0.6 was reached. The temperature was reduced to 25 °C, and 0.15 mM IPTG was added to induce protein expression at an OD_600_ of ∼0.8. Cells were harvested by centrifugation 16 h after induction. All purification steps were performed at 4 °C. Bacterial pellets were resuspended in lysis buffer (50 mM Tris pH 7.5, 300 mM NaCl, 5% glycerol, 10 mM imidazole, 0.1% Triton X-100, 1 mM MgSO_4_). Lysis buffer was supplemented with protease inhibitors (Xpert Protease Inhibitor Cocktail, GenDEPOT), lysozyme, and Deoxyribonuclease I (Worthington). The resuspended pellet was lysed by sonication on ice and cell debris was removed by centrifugation for 25 min at 15,000 *g*. Clarified lysate was loaded into a 20 mL packed IMAC column (Ni-sepharose Excel, Cytiva) and washed with six column volumes of loading buffer (50 mM Tris pH 7.5, 300 mM NaCl, 5% glycerol). G6PD protein was eluted with three column volumes of elution buffer (50 mM Tris pH 7.5, 300 mM NaCl, 400 mM imidazole, 5% glycerol). The eluent was diluted with 20 mM Tris pH 8.0 to a final NaCl concentration of 50 mM. The diluted protein sample was loaded onto a 12 mL Source 15Q anion exchange column (Cytiva) and washed with two column volumes of A buffer (20 mM Tris pH 8.0). Protein was eluted with a linear gradient from 0-50% B buffer (20 mM Tris pH 8.0, 1 M NaCl) over eight column volumes with a fraction size of 2 mL. SDS-PAGE was performed on the eluted fractions to identify the peak corresponding to G6PD. Fractions containing G6PD were pooled and concentrated with a 10 kDa MWCO concentrator (Sartoris) prior to a polishing step using a Superdex 75 pg preparative size-exclusion chromatography (SEC) column. Buffer for SEC was 20 mM HEPES pH 7.5, 150 mM NaCl. SDS-PAGE was performed on the eluted fractions to identify the peak corresponding to G6PD. The fractions corresponding to pure G6PD were pooled and concentrated using a 10 kDa MWCO concentrator (Sartoris). Final G6PD sample purity was evaluated by analytical size-exclusion chromatography using a Superdex 200 10/300 column (Cytiva) and the same SEC buffer. Protein was aliquoted and kept frozen at -80°C.

#### G6PD Activity assay

The enzymatic activity of purified recombinant wild-type (WT), African A⁻ (V68M/N126D), and Mediterranean (S188F) G6PD proteins was determined spectrophotometrically by monitoring the rate of NADPH production via UV absorbance at a wavelength of 340 nm in a Nanodrop 1C (Thermofisher, Waltham, MA). A buffer of 30 mM Tris-HCl (pH 7.8), 2 mM MgCl₂, 69 µM NADP⁺, and 33 mM glucose-6-phosphate was used for this assay which was conducted at 25 °C in a quartz cuvette. G6PD enzyme concentration was 1.44 µM for all measurements. Activity values were expressed as a percentage relative to WT G6PD. To assess the effect of aging at 37 °C, G6PD variants were diluted to a concentration of 0.4 mg/mL in a buffer consisting of 20 mM HEPES (pH 7.5), 100 mM NaCl, 10µM NADP^+^ and placed in an incubator set to a temperature of 37 °C. Samples were periodically sampled (0d, 3d, 5d, 7d) and G6PD activity assayed in the same manner as before. An additional aliquot of each G6PD variant was aged at 4 °C in a refrigerator for 7d. The G6PD activity of the 4 °C samples were only measured at the 7d timepoint. A BCA assay was used to assess soluble G6PD concentration at the 7d timepoint for both the 37 °C and 4 °C aged samples.

#### Analytical ultracentrifugation

Sedimentation velocity experiments were carried out using a Beckman Coulter ProteomeLab XL-I analytical ultracentrifuge equipped with an An-60 Ti rotor. Purified recombinant G6PD variants (0.5-1 mg/mL) were loaded in 50 mM Bis-Tris (pH 7), 150 mM NaCl. Absorbance at 280 nm was monitored during centrifugation at 50,000 rpm at 25 °C. Data were analyzed using SEDFIT (continuous c(s) distribution) to determine sedimentation coefficients and relative populations of monomeric, dimeric, tetrameric and higher-order oligomeric species.

#### Mass photometry

Single-molecule mass distributions were measured on a Refeyn 2MP instrument. Cleaned glass coverslips were assembled into flow chambers, and recombinant proteins were diluted to 1–10 nM in 25 mM HEPES (pH 7.5) w/150 mM NaCl immediately before acquisition. Binding events were recorded at room temperature for 60 s and sample concentrations were adjusted to result in 1000-4000 recorded events. Mass distributions were extracted with Refeyn software to estimate the relative abundance of oligomeric states.

#### Differential Scanning fluorimetry

Thermal stability of G6PD variants was measured by differential scanning fluorimetry using a Bio-Rad 384 OPUS (Bio-Rad, Hercules, CA). G6PD proteins (5 µM in 20 mM HEPES (pH 7.4), 100 mM NaCl) were heated from 25 °C to 95 °C at 1 °C/min in the presence or absence of 0, 10 or 100 µM NADP⁺. SYPRO Orange dye (490 nm excitation, 600 nm emission) was used to monitor protein unfolding and Tm was calculated from the midpoint (first derivative) of the melting transition^111^.

#### Differential Scanning Calorimetry (DSC)

DSC measurements were performed on G6PD wild-type, A- and Med- variants using a Malvern PEAQ-DSC (Malvern Pananalytical, Worcestershire, UK). Protein samples were resuspended in 20 mM HEPES, 100 mM NaCl, 50 µM NADP+ pH 7.4 buffer and protein was exchanged with this buffer for 10 volume equivalents using a spin concentrator column. Sample concentration was determined by UV absorbance at 280 using a Nanodrop One^C^ (Thermofisher, Waltham, MA). Protein at a concentration of 8 µM was placed inside the DSC cell with the buffer used as the reference and was heated from 20 - 75°C at a ramp rate of 60°C/hr inside the DSC instrument. The feedback mode for the experiment was set to low. Data analysis was accomplished using the Malvern PEAQ-DSC software package and the resulting melt curves were fit using a two-transition, non 2-state model providing Tm, ΔH and ΔHVH for each transition.

#### Infrared Spectroscopy

Fourier-transform infrared (FTIR) spectroscopy was performed to assess secondary structure and thermal stability of recombinant G6PD proteins. Purified canonical, A⁻ (V68M/N126D), and Med− (S188F) enzymes were buffer-exchanged into 20 mM HEPES, pH 7.4, 150 mM NaCl, and concentrated to ∼1 mg/mL. Spectra were collected using a JASCO 815 CD spectrometer equipped with an attenuated total reflectance (ATR) module. Samples were scanned over the mid-IR region (1,800–1,400 cm⁻¹) at 1 cm⁻¹ resolution with 128 accumulations per spectrum. Temperature-dependent unfolding was assessed by incrementally heating samples from 25 °C to 90 °C (1 °C/min) with a Peltier-controlled cell holder. Raw spectra were baseline-corrected and vector-normalized prior to analysis. Amide I (1,600–1,700 cm⁻¹) bands were deconvoluted by second-derivative and curve-fitting analysis (OriginPro) to estimate relative α-helical, β-sheet, and random-coil content. Transition midpoints (Ton, Tm) were derived from sigmoidal fits of temperature-dependent band shifts.

#### Negative-Stain Electron Microscopy

Negative-stain transmission electron microscopy (TEM) was used to visualize oligomeric states of recombinant G6PD proteins. Canonical, A⁻, and Med⁻ variants were purified to homogeneity and diluted to 0.05–0.1 mg/mL in 20 mM HEPES, pH 7.4, 150 mM NaCl. Samples (3-5 µL) were adsorbed onto glow-discharged carbon-coated copper grids (400 mesh, EMS) for 1 min, washed briefly with ultrapure water, and stained with 1.5% (w/v) uranyl acetate for 0.5 min. Grids were imaged on a Thermo Fisher Talos L120C transmission electron microscope operated at 120 kV, and micrographs were recorded at a nominal magnification of 73,000× using a Ceta CMOS camera.

### Cryo-EM sample preparation

3 uL of purified G6PD at a concentration of 0.4 mg/mL (A-) or 0.5 mg/mL (WT/Med-) was applied to Quantifoil 1.2/1.3 200 mesh copper grids that were glow discharged for 60s. G6PD samples were plunge-frozen into liquid ethane using the FEI Vitrobot Mark IV (Thermo Fisher Scientific) while maintaining 100% humidity at 4 °C in the blotting chamber. A blot time of 7s, a blot force of 0, and a wait time of 10s was used for G6PD A-. A blot time of 3s, a blot force of -6, and a wait time of 10s was used for G6PD WT and G6PD Med-.

### Cryo-EM data collection

7,679 movies of G6PD A-were recorded on a Krios G3i 300kV TEM equipped with a Falcon4i direct detection camera and Selectris energy filter. A nominal magnification of 130,000x was used, with a pixel size of 0.99 Å, and movies were captured with a defocus range of -0.6 to -1.6. Each movie consisted of 40 frames with a total dose of 50.0 e-/Å^2^. 5,863 and 11,217 movies were captured of G6PD WT and G6PD Med-, respectively, using an Arctica 200kV TEM equipped with a Falcon4i direct detection camera and Selectris energy filter. A nominal magnification of 130,000x was used, with a physical pixel size of 0.899 Å/pixel. A nominal magnification of 130,000x was used, with a pixel size of 0.899 Å, and movies were captured with a defocus range of -0.6 to -1.6. Each movie consisted of 40 frames with a total dose of 50.0 e-/Å^2^.

### Cryo-EM data processing

All movies were imported into CryoSPARC v5.0.4. The Patch Motion Correction tool was used to perform motion correction, and contrast transfer functions were calculated using the Patch CTF Estimation tool. Low-quality micrographs were discarded using the Curate Exposures tool. Particles were extracted with a box size of 240 pixels binned to 60 pixels. 2D class averages were generated to create an ab initio model, which was used as an initial volume for the Heterogeneous Refinement tool. Heterogeneous Refinement and 3D Classification jobs were used to sort particles at increasing resolution according to the pipeline in **Figure S6-7**. Final models were created from non-duplicate, full-resolution particles that underwent Non-Uniform Refinement (D2 Symmetry), Local CTF Refinement, and Reference-Based Motion Correction.

### Model Building and Refinement

Model building and refinement were carried out using an Alphafold2^112^ predicted structures as starting models, which were fit into the corresponding maps using UCSF ChimeraX. Models were refined through multiple rounds of resolve CryoEM and real-space refinement in Phenix v1.21.541957 combined with manual adjustment in COOT (v0.9.8.92).

#### Cross-linking proteomics across a thermal gradient with TMT10

To interrogate structural remodeling of G6PD variants under thermal stress, cross-linking proteomics with tandem mass tag (TMT10) multiplexing was performed. Recombinant proteins (0.5 mg/mL) were incubated with 1 mM di(sulfosuccinimidyl)suberate (DSSO) at 25 °C for 30 min, followed by quenching with 50 mM Tris-HCl (pH 7.5). Cross-linked proteins were subjected to a temperature gradient (25-70 °C, 5 °C increments) using a thermocycler, digested with trypsin, and peptides were labeled with TMT10 reagents according to the manufacturer’s instructions (Thermo Fisher). Labeled peptides were pooled, fractionated by high-pH reversed-phase chromatography, and analyzed by nanoLC-MS/MS on an Orbitrap Fusion Lumos mass spectrometer. Cross-linked peptides were identified using the XlinkX search node in Proteome Discoverer, and relative abundances across the thermal gradient were quantified by TMT reporter ion intensities. Data were used to infer temperature-dependent remodeling of oligomer interfaces and structural elements in WT and mutant G6PD proteins.

### Methemoglobin Measurements

Hemoglobin (Hb) concentration was measured using an ABL90 FLEX blood gas analyzer (Radiometer America, Brea, CA, USA) at 4-minute intervals for the first 20 minutes of incubation and at 5-minute intervals for the last 40 minutes of incubation with 150 µM 5,6-primaquine orthoquinone, as described.^88^ The rates of methemoglobin (MetHb) generation were calculated as the ratio of MetHb produced between successive measurements to the time interval between measurements.

#### Measurement of superoxide in RBCs by Electron Paramagnetic Resonance (EPR) spectroscopy

All EPR spectra were recorded on a spectrometer operating at X-band (9.65 GHz; EMXnano, Bruker Corp., Billerica, MA). RBCs were stored on ice until immediately before measurement. For measuring superoxide, each sample was made by mixing 2 μL of 50 mM CMH stock solution with 98 μL of RBC suspension (40% hematocrit) in PBS-G. Time zero (t = 0) for each measurement is the time at which CMH was mixed into the sample. Approximately 50 μL of each sample was loaded into a 50-μL calibrated borosilicate capillary micropipette (Drummond Scientific Company, Broomall, PA) whose ends were then closed with sealing compound (Chā-seal, DWK Life Sciences, Millville, NJ). The sealed micropipette was positioned in the resonator of the spectrometer for spectral acquisition. The amplitude (h) of the downfield (leftmost) peak of the CM• nitroxide EPR spectrum was converted to concentration (C in μM) by using the calibration equation C = (h – 0.03308)/0.06461, determined using a series of standard samples. A linear least-squares fit of the C-vs-t plot yields a slope that is the rate of CMH oxidation by the RBC sample; the slope is directly proportional to the mean steady-state superoxide concentration in the RBCs. For each type of RBC (B6, hG6PD, A-, Med-), at least four replicate measurements were performed.

## Acknowledgments

The content is solely the responsibility of the authors and does not represent the official views of the National Institutes of Health. The authors would like to thank all the donor volunteers who participated in this study and all the global blood donor communities for their life-saving altruistic gifts. We thank Drs. John Hardin of the CU Anschutz cryo-EM facility for assistance with EM data collection. Cryo-EM data collection for the A- tetramer was collected at the Biochemistry Krios Electron Microscopy Facility (BioKEM) at CU Boulder (RRID: SCR_019057).

## Author Contribution

Clinical trial in SCD: MSK, SLS; Animal studies: AH, MSP, EAL, DRL, PWB, JCZ. Proteomics: MD, KCH. Metabolomics and lipidomics analyses: JAR, DS, FC, TN, AD. Biostatistics and Bioinformatics: GRK, ALM, XD, GPP, NR, AD. REDS RBC Omics: MS, SK, SLS, PJN, MPB. Structural Studies and Biochemistry: SB, AVI, EM, RRS, FV, JPYK, EWH, JJ, EZE, PWB; Systems biology models: ZBH, AMK, BOP; Vein-to-vein database: NHR; mQTL and pQTL analyses: GRK, ALM, GPP. Figure preparation: RRS, EM, JJ, AD, GRK, PWB. Conceptualization: YX, JCZ, AD. Supervision: AD, JJ, KCH, PJN, MPB, BOP, SLS, GPP, JCZ, PWB. Writing and finalization: first draft by AD and all co-authors reviewed and approved the final version.

## Clinical trial

The NHLBI Recipient Epidemiology Donor Evaluation Study (REDS)-III Red Blood Cell Omics (RBC-Omics) and Vein to Vein databases are accessible at https://biolincc.nhlbi.nih.gov/studies/reds_iii/ and Genomics data are deposited at dbGaP Study Accession: phs001955.v1.p1 “The Impact of Oxidative Stress on Erythrocyte Biology (RBC Survival)” trial is accessible through ClinicalTrials.gov ID: NCT04028700.

## Data and Materials availability

Accession PDB numbers for the deposited structures are listed below:

- WT tetramer with NADP: 39FH
- Structure of A- G6PD (V68M,N126D) tetramer with NADP: 39FT
- Structure of Med (S188F) G6PD tetramer with NADP: 39FV
- Structure of Med (S188F) G6PD tetramer bound to NADP-3DVA - 39FW
- Structure of Med (S188F) G6PD dimer bound to NADP: 39FX

The proteomics datasets are available through MassIVE with identifier MSV000103263. The metabolomics data are available through Metabolomics Workbench with study identifier ST005223. All raw data and elaborations are included in Supplementary Table 1.xlsx. Humanized G6PD canonical or deficient mice (African and Mediterranean variants) are available upon reasonable request, finalization of material transfer agreement and after institutional ACUC approval through Dr James C Zimring Lab. All three-dimensional cryo-EM density maps, coordinates for the atomic models and local-refined maps generated in this study have been deposited and are publicly available as of the date of publication. Further information and requests for resources and reagents should be directed to and will be fulfilled by the Lead Contact, Angelo D’Alessandro.

## REFERENCES

1. Nkhoma, E.T., Poole, C., Vannappagari, V., Hall, S.A., and Beutler, E. (2009). The global prevalence of glucose-6-phosphate dehydrogenase deficiency: a systematic review and meta-analysis. Blood Cells Mol Dis 42, 267–278. 10.1016/j.bcmd.2008.12.005.

2. Luzzatto, L., Bancone, G., Dugué, P.A., Jiang, W., Minucci, A., Nannelli, C., Pfeffer, D., Prchal, J., Sirdah, M., Sodeinde, O., et al. (2024). New WHO classification of genetic variants causing G6PD deficiency. Bull World Health Organ 102, 615–617. 10.2471/blt.23.291224.

3. TeSlaa, T., Ralser, M., Fan, J., and Rabinowitz, J.D. (2023). The pentose phosphate pathway in health and disease. Nat Metab 5, 1275–1289. 10.1038/s42255-023-00863-2.

4. Chatzinikolaou, P.N., Margaritelis, N.V., Paschalis, V., Theodorou, A.A., Vrabas, I.S., Kyparos, A., D’Alessandro, A., and Nikolaidis, M.G. (2024). Erythrocyte metabolism. Acta Physiologica 240, e14081. 10.1111/apha.14081.

5. Au, S.W., Gover, S., Lam, V.M., and Adams, M.J. (2000). Human glucose-6-phosphate dehydrogenase: the crystal structure reveals a structural NADP(+) molecule and provides insights into enzyme deficiency. Structure 8, 293–303. 10.1016/s0969-2126(00)00104-0.

6. Horikoshi, N., Hwang, S., Gati, C., Matsui, T., Castillo-Orellana, C., Raub, A.G., Garcia, A.A., Jabbarpour, F., Batyuk, A., Broweleit, J., et al. (2021). Long-range structural defects by pathogenic mutations in most severe glucose-6-phosphate dehydrogenase deficiency. Proceedings of the National Academy of Sciences 118, e2022790118. doi:10.1073/pnas.2022790118.

7. Hwang, S., Mruk, K., Rahighi, S., Raub, A.G., Chen, C.-H., Dorn, L.E., Horikoshi, N., Wakatsuki, S., Chen, J.K., and Mochly-Rosen, D. (2018). Correcting glucose-6-phosphate dehydrogenase deficiency with a small-molecule activator. Nature Communications 9, 4045. 10.1038/s41467-018-06447-z.

8. Hanau, S., and Helliwell, J.R. (2024). Glucose-6-phosphate dehydrogenase and its 3D structures from crystallography and electron cryo-microscopy. Acta Crystallogr F Struct Biol Commun 80, 236–251. 10.1107/s2053230x24008112.

9. Yoshida, T., Prudent, M., and D’Alessandro, A. (2019). Red blood cell storage lesion: causes and potential clinical consequences. Blood Transfus 17, 27–52. 10.2450/2019.0217-18.

10. D’Alessandro, A., Keele, G.R., Hay, A., Nemkov, T., Earley, E.J., Stephenson, D., Vincent, M., Deng, X., Stone, M., Dzieciatkowska, M., et al. (2025). Ferroptosis regulates hemolysis in stored murine and human red blood cells. Blood 145, 765–783. 10.1182/blood.2024026109.

11. Reisz, J.A., Wither, M.J., Dzieciatkowska, M., Nemkov, T., Issaian, A., Yoshida, T., Dunham, A.J., Hill, R.C., Hansen, K.C., and D’Alessandro, A. (2016). Oxidative modifications of glyceraldehyde 3-phosphate dehydrogenase regulate metabolic reprogramming of stored red blood cells. Blood 128, e32–42. 10.1182/blood-2016-05-714816.

12. Rogers, S.C., Ge, X., Brummet, M., Lin, X., Timm, D.D., d’Avignon, A., Garbow, J.R., Kao, J., Prakash, J., Issaian, A., et al. (2021). Quantifying dynamic range in red blood cell energetics: Evidence of progressive energy failure during storage. Transfusion 61, 1586–1599. 10.1111/trf.16395.

13. Peltier, S., Marin, M., Dzieciatkowska, M., Dussiot, M., Roy, M.K., Bruce, J., Leblanc, L., Hadjou, Y., Georgeault, S., Fricot, A., et al. (2025). Proteostasis and metabolic dysfunction characterize a subset of storage-induced senescent erythrocytes targeted for posttransfusion clearance. J Clin Invest 135. 10.1172/jci183099.

14. Messana, I., Ferroni, L., Misiti, F., Girelli, G., Pupella, S., Castagnola, M., Zappacosta, B., and Giardina, B. (2000). Blood bank conditions and RBCs: the progressive loss of metabolic modulation. Transfusion 40, 353–360. 10.1046/j.1537-2995.2000.40030353.x.

15. Luzzatto, L., Ally, M., and Notaro, R. (2020). Glucose-6-phosphate dehydrogenase deficiency. Blood 136, 1225–1240. 10.1182/blood.2019000944.

16. Francis, R.O., Jhang, J.S., Pham, H.P., Hod, E.A., Zimring, J.C., and Spitalnik, S.L. (2013). Glucose-6-phosphate dehydrogenase deficiency in transfusion medicine: the unknown risks. Vox Sanguinis 105, 271–282. 10.1111/vox.12068.

17. Francis, R.O., D’Alessandro, A., Eisenberger, A., Soffing, M., Yeh, R., Coronel, E., Sheikh, A., Rapido, F., La Carpia, F., Reisz, J.A., et al. (2020). Donor glucose-6-phosphate dehydrogenase deficiency decreases blood quality for transfusion. J Clin Invest 130, 2270–2285. 10.1172/jci133530.

18. D’Alessandro, A., Fu, X., Kanias, T., Reisz, J.A., Culp-Hill, R., Guo, Y., Gladwin, M.T., Page, G., Kleinman, S., Lanteri, M., et al. (2021). Donor sex, age and ethnicity impact stored red blood cell antioxidant metabolism through mechanisms in part explained by glucose 6-phosphate dehydrogenase levels and activity. Haematologica 106, 1290–1302. 10.3324/haematol.2020.246603.

19. Sagiv, E., Fasano, R.M., Luban, N.L.C., Josephson, C.D., Stowell, S.R., Roback, J.D., Francis, R.O., and Yee, M.E.M. (2018). Glucose-6-phosphate-dehydrogenase deficient red blood cell units are associated with decreased posttransfusion red blood cell survival in children with sickle cell disease. Am J Hematol 93, 630–634. 10.1002/ajh.25051.

20. Yee, M.E., Francis, R.O., Luban, N.L.C., Easley, K.A., Lough, C.M., Roback, J.D., Josephson, C.D., and Fasano, R.M. (2022). Glucose-6-phosphate dehydrogenase deficiency is more prevalent in Duffy-null red blood cell transfusion in sickle cell disease. Transfusion 62, 551–555. 10.1111/trf.16806.

21. Le Gallo, M., Moutereau, S., Gentil, M., and Pirenne, F. (2024). Study of the antigenic characteristics of red blood cells units and their sickle cell disease recipients and the G6PD activity of transfused red blood cells units. Transfus Clin Biol 31, 130–134. 10.1016/j.tracli.2024.03.002.

22. Maffi, D., Pasquino, M.T., Mandarino, L., Tortora, P., Girelli, G., Meo, D., Grazzini, G., and Caprari, P. (2014). Glucose-6-phosphate dehydrogenase deficiency in Italian blood donors: prevalence and molecular defect characterization. Vox Sang 106, 227–233. 10.1111/vox.12096.

23. La Raja, M., La Rocca, U., Forni, G.L., Garozzo, G., Manca, P.C., Pupella, S., and Teofili, L. Eligibility of blood donors with thalassaemia trait, sickle cell trait or glucose-6-phosphate-dehydrogenase deficiency in Italy: Results of a national survey and practical guidance. Vox Sanguinis n/a. 10.1111/vox.70361.

24. Page, G.P., Kanias, T., Guo, Y.J., Lanteri, M.C., Zhang, X., Mast, A.E., Cable, R.G., Spencer, B.R., Kiss, J.E., Fang, F., et al. (2021). Multiple-ancestry genome-wide association study identifies 27 loci associated with measures of hemolysis following blood storage. J Clin Invest 131. 10.1172/jci146077.

25. Roubinian, N.H., Reese, S.E., Qiao, H., Plimier, C., Fang, F., Page, G.P., Cable, R.G., Custer, B., Gladwin, M.T., Goel, R., et al. (2022). Donor genetic and nongenetic factors affecting red blood cell transfusion effectiveness. JCI Insight 7. 10.1172/jci.insight.152598.

26. Tzounakas, V.L., Kriebardis, A.G., Georgatzakou, H.T., Foudoulaki-Paparizos, L.E., Dzieciatkowska, M., Wither, M.J., Nemkov, T., Hansen, K.C., Papassideri, I.S., D’Alessandro, A., and Antonelou, M.H. (2016). Glucose 6-phosphate dehydrogenase deficient subjects may be better “storers” than donors of red blood cells. Free Radic Biol Med 96, 152–165. 10.1016/j.freeradbiomed.2016.04.005.

27. Josephson, C.D., Glynn, S., Mathew, S., Birch, R., Bakkour, S., Baumann Kreuziger, L., Busch, M.P., Chapman, K., Dinardo, C., Hendrickson, J., et al. (2022). The Recipient Epidemiology and Donor Evaluation Study-IV-Pediatric (REDS-IV-P): A research program striving to improve blood donor safety and optimize transfusion outcomes across the lifespan. Transfusion 62, 982–999. 10.1111/trf.16869.

28. D’Alessandro, A., Howie, H.L., Hay, A.M., Dziewulska, K.H., Brown, B.C., Wither, M.J., Karafin, M., Stone, E.F., Spitalnik, S.L., Hod, E.A., et al. (2021). Hematologic and systemic metabolic alterations due to Mediterranean class II G6PD deficiency in mice. JCI Insight 6. 10.1172/jci.insight.147056.

29. Wang, L., Rochon, E.R., Gingras, S., Zuchelkowski, B.E., Sinchar, D.J., Alipour, E., Reisz, J.A., Yang, M., Page, G.P., Kanias, T., et al. (2024). Functional effects of an African glucose-6-phosphate dehydrogenase (G6PD) polymorphism (Val68Met) on red blood cell hemolytic propensity and post-transfusion recovery. Transfusion 64, 615–626. 10.1111/trf.17756.

30. Baek, J.H., Yalamanoglu, A., Moon, S.-E., Gao, Y., and Buehler, P.W. (2018). Evaluation of renal oxygen homeostasis in a preclinical animal model to elucidate difference in blood quality after transfusion. Transfusion 58, 1474–1485. 10.1111/trf.14560.

31. Dumbill, R., Rabcuka, J., Fallon, J., Knight, S., Hunter, J., Voyce, D., Barrett, J., Ellen, M., Weissenbacher, A., Kurniawan, T., et al. (2024). Impaired O2 unloading from stored blood results in diffusion-limited O2 release at tissues: evidence from human kidneys. Blood 143, 721–733. 10.1182/blood.2023022385.

32. Wei, X., Kixmoeller, K., Baltrusaitis, E., Yang, X., and Marmorstein, R. (2022). Allosteric role of a structural NADP^+^ molecule in glucose-6-phosphate dehydrogenase activity. Proceedings of the National Academy of Sciences 119, e2119695119. doi:10.1073/pnas.2119695119.

33. Andrews, P.H., Zimring, J.C., and McNamara, C.A. (2025). Clinical associations and potential cellular mechanisms linking G6PD deficiency and atherosclerotic cardiovascular disease. npj Metabolic Health and Disease 3, 16. 10.1038/s44324-025-00061-6.

34. Nemkov, T., Key, A., Stephenson, D., Earley, E.J., Keele, G.R., Hay, A.M., Amireault, P., Casimir, M., Dussiot, M., Dzieciatkowska, M., et al. (2024). Genetic regulation of carnitine metabolism controls lipid damage repair and aging RBC hemolysis in vivo and in vitro. Blood. 10.1182/blood.2024023983.

35. Nemkov, T., Sun, K., Reisz, J.A., Yoshida, T., Dunham, A., Wen, E.Y., Wen, A.Q., Roach, R.C., Hansen, K.C., Xia, Y., and D’Alessandro, A. (2017). Metabolism of Citrate and Other Carboxylic Acids in Erythrocytes As a Function of Oxygen Saturation and Refrigerated Storage. Front Med (Lausanne) 4, 175. 10.3389/fmed.2017.00175.

36. Bordbar, A., Yurkovich, J.T., Paglia, G., Rolfsson, O., Sigurjónsson Ó, E., and Palsson, B.O. (2017). Elucidating dynamic metabolic physiology through network integration of quantitative time-course metabolomics. Sci Rep 7, 46249. 10.1038/srep46249.

37. Maurya, P.K., Kumar, P., and Chandra, P. (2016). Age-dependent detection of erythrocytes glucose-6-phosphate dehydrogenase and its correlation with oxidative stress. Archives of Physiology and Biochemistry 122, 61–66. 10.3109/13813455.2015.1136648.

38. van den Hurk, K., Zalpuri, S., Prinsze, F.J., Merz, E.-M., and de Kort, W.L.A.M. (2017). Associations of health status with subsequent blood donor behavior—An alternative perspective on the Healthy Donor Effect from Donor InSight. PLOS ONE 12, e0186662. 10.1371/journal.pone.0186662.

39. He, F., Ru, X., and Wen, T. (2020). NRF2, a Transcription Factor for Stress Response and Beyond. Int J Mol Sci 21. 10.3390/ijms21134777.

40. Hay, A.M., Nemkov, T., Gamboni, F., Dzieciatkowska, M., Key, A., Galbraith, M., Bartsch, K., Sun, K., Xia, Y., Stone, M., et al. (2022). S1P has a negative effect on RBC storage quality. Blood Adv. 10.1182/bloodadvances.2022008936.

41. Karafin, M.S., Grier, A.L., Fasano, R.M., Ilich, A., Wichlan, D., Chang, A., James, S.M., Butler, H.E., Kolupaev, O., Caughey, M.C., et al. (2025). Blood-storage duration affects hematological and metabolic profiles in patients with sickle cell disease receiving transfusions. J Clin Invest 135. 10.1172/jci192920.

42. Roubinian, N.H., Kleinman, S., Murphy, E.L., Glynn, S.A., and Edgren, G. (2020). Methodological considerations for linked blood donor-component-recipient analyses in transfusion medicine research. ISBT Science Series 15, 185–193. 10.1111/voxs.12518.

43. Unali, G., Crivicich, G., Pagani, I., Abou-Alezz, M., Folchini, F., Valeri, E., Matafora, V., Reisz, J.A., Giordano, A.M.S., Cuccovillo, I., et al. (2023). Interferon-inducible phospholipids govern IFITM3-dependent endosomal antiviral immunity. Embo j 42, e112234. 10.15252/embj.2022112234.

44. Xu, P., Chen, C., Zhang, Y., Dzieciatkowska, M., Brown, B.C., Zhang, W., Xie, T., Abdulmalik, O., Song, A., Tong, C., et al. (2022). Erythrocyte transglutaminase-2 combats hypoxia and chronic kidney disease by promoting oxygen delivery and carnitine homeostasis. Cell Metab 34, 299–316.e296. 10.1016/j.cmet.2021.12.019.

45. Sun, K., Zhang, Y., D’Alessandro, A., Nemkov, T., Song, A., Wu, H., Liu, H., Adebiyi, M., Huang, A., Wen, Y.E., et al. (2016). Sphingosine-1-phosphate promotes erythrocyte glycolysis and oxygen release for adaptation to high-altitude hypoxia. Nat Commun 7, 12086. 10.1038/ncomms12086.

46. D’Alessandro, A., Earley, E.J., Nemkov, T., Stephenson, D., Dzieciatkowska, M., Hansen, K.C., Minetti, G., Champigneulle, B., Stauffer, E., Pichon, A., et al. (2024). Genetic polymorphisms and expression of Rhesus blood group RHCE are associated with 2,3-bisphosphoglycerate in humans at high altitude. Proc Natl Acad Sci U S A 121, e2315930120. 10.1073/pnas.2315930120.

47. D’Alessandro, A., Blasi, B., D’Amici, G.M., Marrocco, C., and Zolla, L. (2013). Red blood cell subpopulations in freshly drawn blood: application of proteomics and metabolomics to a decades-long biological issue. Blood Transfus 11, 75–87. 10.2450/2012.0164-11.

48. Piomelli, S., Corash, L.M., Davenport, D.D., Miraglia, J., and Amorosi, E.L. (1968). In vivo lability of glucose-6-phosphate dehydrogenase in GdA- and GdMediterranean deficiency. J Clin Invest 47, 940–948. 10.1172/jci105786.

49. Cancedda, R., Ogunmola, G., and Luzzatto, L. (1973). Genetic variants of human erythrocyte glucose-6-phosphate dehydrogenase. Discrete conformational states stabilized by NADP + and NADPH. Eur J Biochem 34, 199–204. 10.1111/j.1432-1033.1973.tb02746.x.

50. Ramírez-Nava, E.J., Ortega-Cuellar, D., Serrano-Posada, H., González-Valdez, A., Vanoye-Carlo, A., Hernández-Ochoa, B., Sierra-Palacios, E., Hernández-Pineda, J., Rodríguez-Bustamante, E., Arreguin-Espinosa, R., et al. (2017). Biochemical Analysis of Two Single Mutants that Give Rise to a Polymorphic G6PD A-Double Mutant. Int J Mol Sci 18. 10.3390/ijms18112244.

51. Punjani, A., and Fleet, D.J. (2021). 3D variability analysis: Resolving continuous flexibility and discrete heterogeneity from single particle cryo-EM. J Struct Biol 213, 107702. 10.1016/j.jsb.2021.107702.

52. Nagel, Z.D., and Klinman, J.P. (2010). Update 1 of: Tunneling and dynamics in enzymatic hydride transfer. Chem Rev 110, Pr41–67. 10.1021/cr1001035.

53. Reisz, J.A., Nemkov, T., Dzieciatkowska, M., Culp-Hill, R., Stefanoni, D., Hill, R.C., Yoshida, T., Dunham, A., Kanias, T., Dumont, L.J., et al. (2018). Methylation of protein aspartates and deamidated asparagines as a function of blood bank storage and oxidative stress in human red blood cells. Transfusion 58, 2978–2991. 10.1111/trf.14936.

54. Bevers, S., Issaian, A.V., Hay, A., Keele, G.R., Dzieciatkowska, M., Reisz, J.A., Haiman, Z.B., Nemkov, T., Stephenson, D., Moore, A.L., et al. (2026). Improved red blood cell storage quality of blood from donors carrying the hypermorphic PIMT I120 variant. Hemasphere 10, e70361. 10.1002/hem3.70361.

55. Stephenson, D., Keele, G.R., Hay, A., Dzieciatkowska, M., Reisz, J.A., Haiman, Z.B., Moore, A.L., Nemkov, T., Deng, X., Stone, M., et al. (2025). GPX4 regulates lipid peroxidation and ferroptosis of stored red blood cells. Blood Red Cells & Iron, 100020. 10.1016/j.brci.2025.100020.

56. Kanias, T., Lanteri, M.C., Page, G.P., Guo, Y., Endres, S.M., Stone, M., Keating, S., Mast, A.E., Cable, R.G., Triulzi, D.J., et al. (2017). Ethnicity, sex, and age are determinants of red blood cell storage and stress hemolysis: results of the REDS-III RBC-Omics study. Blood Adv 1, 1132–1141. 10.1182/bloodadvances.2017004820.

57. Babalola, A.O., Beetlestone, J.G., and Luzzatto, L. (1976). Genetic variants of human erythrocyte glucose-6-phosphate dehydrogenase. Kinetic and thermodynamic parameters of variants A, B, and A- in relation to quaternary structure. Journal of Biological Chemistry 251, 2993–3002. 10.1016/S0021-9258(17)33490-7.

58. Kotaka, M., Gover, S., Vandeputte-Rutten, L., Au, S.W., Lam, V.M., and Adams, M.J. (2005). Structural studies of glucose-6-phosphate and NADP+ binding to human glucose-6-phosphate dehydrogenase. Acta Crystallogr D Biol Crystallogr 61, 495–504. 10.1107/s0907444905002350.

59. Batista da Rocha, J., Othman, H., and Hazelhurst, S. (2022). Molecular dynamics of G6PD variants from sub-Saharan Africa. Biochemistry and Biophysics Reports 30, 101236. 10.1016/j.bbrep.2022.101236.

60. Garcia, A.A., Mathews, II, Horikoshi, N., Matsui, T., Kaur, M., Wakatsuki, S., and Mochly-Rosen, D. (2022). Stabilization of glucose-6-phosphate dehydrogenase oligomers enhances catalytic activity and stability of clinical variants. J Biol Chem 298, 101610. 10.1016/j.jbc.2022.101610.

61. Roussel, C., Morel, A., Dussiot, M., Marin, M., Colard, M., Fricot-Monsinjon, A., Martinez, A., Chambrion, C., Henry, B., Casimir, M., et al. (2021). Rapid clearance of storage-induced microerythrocytes alters transfusion recovery. Blood 137, 2285–2298. 10.1182/blood.2020008563.

62. Vallelian, F., Buehler, P.W., and Schaer, D.J. (2022). Hemolysis, free hemoglobin toxicity, and scavenger protein therapeutics. Blood 140, 1837–1844. 10.1182/blood.2022015596.

63. D’Alessandro, A., Nouraie, S.M., Zhang, Y., Cendali, F., Gamboni, F., Reisz, J.A., Zhang, X., Bartsch, K.W., Galbraith, M.D., Espinosa, J.M., Gordeuk, V.R., and Gladwin, M.T. (2023). Metabolic signatures of cardiorenal dysfunction in plasma from sickle cell patients as a function of therapeutic transfusion and hydroxyurea treatment. Haematologica. 10.3324/haematol.2023.283288.

64. Nemkov, T., Stefanoni, D., Bordbar, A., Issaian, A., Palsson, B.O., Dumont, L.J., Hay, A.M., Song, A., Xia, Y., Redzic, J.S., et al. (2020). Blood donor exposome and impact of common drugs on red blood cell metabolism. JCI Insight. 10.1172/jci.insight.146175.

65. Cappelletti, S., Piacentino, D., Sani, G., and Aromatario, M. (2015). Caffeine: cognitive and physical performance enhancer or psychoactive drug? Curr Neuropharmacol 13, 71–88. 10.2174/1570159x13666141210215655.

66. Xu, H., Hu, L., Liu, T., Chen, F., Li, J., Xu, J., Jiang, L., Xiang, Z., Wang, X., and Sheng, J. (2020). Caffeine Targets G6PDH to Disrupt Redox Homeostasis and Inhibit Renal Cell Carcinoma Proliferation. Front Cell Dev Biol 8, 556162. 10.3389/fcell.2020.556162.

67. Dzieciatkowska, M., Hay, A., Issaian, A., Keele, G.R., Bevers, S., Nemkov, T., Reisz, J.A., Maslanka, M., Stephenson, D., Moore, A.L., et al. (2026). Caffeine impairs red blood cell storage quality by dual inhibition of ADORA2b signaling and G6PD activity. Haematologica 111, 861–875. 10.3324/haematol.2025.288332.

68. Gleadall, N.S., Koets, L., Shamardina, O., Gollub, J., Gottschalk, A.J., Razeghi, O., Ochoa-Garay, G., Stephens, J., Varma, R., Martin, J., et al. (2025). Array genotyping of transfusion-relevant blood cell antigens in 6946 ancestrally diverse study participants. Blood 146, 1511–1524. 10.1182/blood.2025028902.

69. Guo, Y., Busch, M.P., Seielstad, M., Endres-Dighe, S., Westhoff, C.M., Keating, B., Hoppe, C., Bordbar, A., Custer, B., Butterworth, A.S., et al. (2019). Development and evaluation of a transfusion medicine genome wide genotyping array. Transfusion 59, 101–111. 10.1111/trf.15012.

70. D’Alessandro, A. (2026). Precision transfusion medicine in the omics era. Blood 148, 1233–1239. 10.1182/blood.2026034666.

71. Jelani, I., Abdullahi, H.L., Alhassan, A.J., Dalhatu, M.M., Dangambo, M.A., Abacha, F.Z., Haruna, S., Mahmud, R.I., Hamza, H.D., Abdu, R., and Wali, U. (2024). Prevalence of Glucose-6-Phosphate Dehydrogenase deficiency among Hausa and Fulani ethnics in Kano, North-West Nigeria. Medical Research Archives 12. 10.18103/mra.v12i5.5314.

72. Endres-Dighe, S.M., Guo, Y., Kanias, T., Lanteri, M., Stone, M., Spencer, B., Cable, R.G., Kiss, J.E., Kleinman, S., Gladwin, M.T., et al. (2019). Blood, sweat, and tears: Red Blood Cell-Omics study objectives, design, and recruitment activities. Transfusion 59, 46–56. 10.1111/trf.14971.

73. D’Alessandro, A., Culp-Hill, R., Reisz, J.A., Anderson, M., Fu, X., Nemkov, T., Gehrke, S., Zheng, C., Kanias, T., Guo, Y., et al. (2019). Heterogeneity of blood processing and storage additives in different centers impacts stored red blood cell metabolism as much as storage time: lessons from REDS-III-Omics. Transfusion 59, 89–100. 10.1111/trf.14979.

74. Nemkov, T., Yoshida, T., Nikulina, M., and D’Alessandro, A. (2022). High-Throughput Metabolomics Platform for the Rapid Data-Driven Development of Novel Additive Solutions for Blood Storage. Frontiers in Physiology 13. 10.3389/fphys.2022.833242.

75. Thomas, T., Stefanoni, D., Dzieciatkowska, M., Issaian, A., Nemkov, T., Hill, R.C., Francis, R.O., Hudson, K.E., Buehler, P.W., Zimring, J.C., et al. (2020). Evidence for structural protein damage and membrane lipid remodeling in red blood cells from COVID-19 patients. medRxiv. 10.1101/2020.06.29.20142703.

76. Reisz, J.A., Zheng, C., D’Alessandro, A., and Nemkov, T. (2019). Untargeted and Semi-targeted Lipid Analysis of Biological Samples Using Mass Spectrometry-Based Metabolomics. Methods Mol Biol 1978, 121–135. 10.1007/978-1-4939-9236-2_8.

77. Stephenson, D., Nemkov, T., Qadri, S.M., Sheffield, W.P., and D’Alessandro, A. (2022). Inductively-Coupled Plasma Mass Spectrometry-Novel Insights From an Old Technology Into Stressed Red Blood Cell Physiology. Front Physiol 13, 828087. 10.3389/fphys.2022.828087.

78. Moore, A., Busch, M.P., Dziewulska, K., Francis, R.O., Hod, E.A., Zimring, J.C., D’Alessandro, A., and Page, G.P. (2022). Genome-wide metabolite quantitative trait loci analysis (mQTL) in red blood cells from volunteer blood donors. J Biol Chem 298, 102706. 10.1016/j.jbc.2022.102706.

79. Nemkov, T., Stephenson, D., Earley, E.J., Keele, G.R., Hay, A., Key, A., Haiman, Z.B., Erickson, C., Dzieciatkowska, M., Reisz, J.A., et al. (2024). Biological and genetic determinants of glycolysis: Phosphofructokinase isoforms boost energy status of stored red blood cells and transfusion outcomes. Cell Metab. 10.1016/j.cmet.2024.06.007.

80. Delaneau, O., Coulonges, C., and Zagury, J.-F. (2008). Shape-IT: new rapid and accurate algorithm for haplotype inference. BMC Bioinformatics 9, 540. 10.1186/1471-2105-9-540.

81. Howie, B., Marchini, J., and Stephens, M. (2011). Genotype Imputation with Thousands of Genomes. G3 Genes|Genomes|Genetics 1, 457–470. 10.1534/g3.111.001198.

82. Zheng, X., Levine, D., Shen, J., Gogarten, S.M., Laurie, C., and Weir, B.S. (2012). A high-performance computing toolset for relatedness and principal component analysis of SNP data. Bioinformatics 28, 3326–3328. 10.1093/bioinformatics/bts606.

83. Aulchenko, Y.S., Struchalin, M.V., and van Duijn, C.M. (2010). ProbABEL package for genome-wide association analysis of imputed data. BMC Bioinformatics 11, 134. 10.1186/1471-2105-11-134.

84. Perry, J.A., Gaynor, B.J., Mitchell, B.D., and O’Connell, J.R. (2021). An Omics Analysis Search and Information System (OASIS) for Enabling Biological Discovery in the Old Order Amish. bioRxiv, 2021.2005.2002.442370. 10.1101/2021.05.02.442370.

85. https://biolincc.nhlbi.nih.gov/studies/reds_iii/.

86. Karafin, M.S., Bruhn, R., Westlake, M., Sullivan, M.T., Bialkowski, W., Edgren, G., Roubinian, N.H., Hauser, R.G., Kor, D.J., Fleischmann, D., et al. (2017). Demographic and epidemiologic characterization of transfusion recipients from four US regions: evidence from the REDS-III recipient database. Transfusion 57, 2903–2913. 10.1111/trf.14370.

87. Endres-Dighe, S.M., Guo, Y., Kanias, T., Lanteri, M., Stone, M., Spencer, B., Cable, R.G., Kiss, J.E., Kleinman, S., Gladwin, M.T., et al. (2019). Blood, sweat, and tears: Red Blood Cell-Omics study objectives, design, and recruitment activities. Transfusion 59, 46–56. 10.1111/trf.14971.

88. Dziewulska-Cronk, K.H., Reisz, J.A., Hay, A.M., Nemkov, T., Cendali, F.I., Issaian, A., Lamb, D.R., Palha, M.S., Legenzov, E.A., Kao, J.P.Y., et al. (2024). Primaquine-5,6-Orthoquinone Is Directly Hemolytic to Older G6PD Deficient RBCs in a Humanized Mouse Model. J Pharmacol Exp Ther 391, 119–129. 10.1124/jpet.124.002218.

89. Cendali, F.I., Grier, A.L., Lisk, C., Dzieciatkowska, M., Haiman, Z., Reisz, J.A., Harral, J., Stephenson, D., Hay, A.M., Wartchow, E.P., et al. (2025). Increased Exercise Tolerance in G6PD African Variant Mice Driven by Metabolic Adaptations and Erythrophagocytosis. Antioxidants (Basel) 14. 10.3390/antiox14080927.

90. Cendali, F.I., Lisk, C., Dzieciatkowska, M., LaCroix, I.S., Reisz, J.A., Harral, J., Stephenson, D., Hay, A.M., Wartchow, E.P., Darehshouri, A., et al. (2025). Increased exercise tolerance in humanized G6PD-deficient mice. Blood Adv 9, 321–334. 10.1182/bloodadvances.2024013968.

91. Hay, A., Dziewulska, K., Gamboni, F., Nerguizian, D., Dzieciatkowska, M., Zimring, J.C., and D’Alessandro, A. (2022). Hypoxic storage of murine red blood cells improves energy metabolism and post-transfusion recoveries. Blood Transfus. 10.2450/2022.0172-22.

92. Hoffmann-Fezer, G., Maschke, H., Zeitler, H.J., Gais, P., Heger, W., Ellwart, J., and Thierfelder, S. (1991). Direct in vivo biotinylation of erythrocytes as an assay for red cell survival studies. Annals of Hematology 63, 214–217. 10.1007/BF01703446.

93. Zimring, J.C., Smith, N., Stowell, S.R., Johnsen, J.M., Bell, L.N., Francis, R.O., Hod, E.A., Hendrickson, J.E., Roback, J.D., and Spitalnik, S.L. (2014). Strain-specific red blood cell storage, metabolism, and eicosanoid generation in a mouse model. Transfusion 54, 137–148. 10.1111/trf.12264.

94. Nemkov, T., Yoshida, T., Nikulina, M., and D’Alessandro, A. (2022). High-Throughput Metabolomics Platform for the Rapid Data-Driven Development of Novel Additive Solutions for Blood Storage. Front Physiol 13, 833242. 10.3389/fphys.2022.833242.

95. Nemkov, T., Reisz, J.A., Gehrke, S., Hansen, K.C., and D’Alessandro, A. (2019). High-Throughput Metabolomics: Isocratic and Gradient Mass Spectrometry-Based Methods. Methods Mol Biol 1978, 13–26. 10.1007/978-1-4939-9236-2_2.

96. Stefanoni, D., Shin, H.K.H., Baek, J.H., Champagne, D.P., Nemkov, T., Thomas, T., Francis, R.O., Zimring, J.C., Yoshida, T., Reisz, J.A., et al. (2020). Red blood cell metabolism in Rhesus macaques and humans: comparative biology of blood storage. Haematologica 105, 2174–2186. 10.3324/haematol.2019.229930.

97. Nemkov, T., Hansen, K.C., and D’Alessandro, A. (2017). A three-minute method for high-throughput quantitative metabolomics and quantitative tracing experiments of central carbon and nitrogen pathways. Rapid Commun Mass Spectrom 31, 663–673. 10.1002/rcm.7834.

98. Nemkov, T., Stephenson, D., Earley, E.J., Keele, G.R., Hay, A., Key, A., Haiman, Z.B., Erickson, C., Dzieciatkowska, M., Reisz, J.A., et al. (2024). Biological and genetic determinants of glycolysis: Phosphofructokinase isoforms boost energy status of stored red blood cells and transfusion outcomes. Cell Metab. 10.1016/j.cmet.2024.06.007.

99. Gautier, E.-F., Leduc, M., Cochet, S., Bailly, K., Lacombe, C., Mohandas, N., Guillonneau, F., El Nemer, W., and Mayeux, P. (2018). Absolute proteome quantification of highly purified populations of circulating reticulocytes and mature erythrocytes. Blood Adv 2, 2646–2657. 10.1182/bloodadvances.2018023515.

100. Bryk, A.H., and Wiśniewski, J.R. (2017). Quantitative Analysis of Human Red Blood Cell Proteome. J. Proteome Res. 16, 2752–2761. 10.1021/acs.jproteome.7b00025.

101. Wiśniewski, J.R., and Rakus, D. (2014). Multi-enzyme digestion FASP and the ’Total Protein Approach’-based absolute quantification of the Escherichia coli proteome. J Proteomics 109, 322–331. 10.1016/j.jprot.2014.07.012.

102. Haiman, Z.B., Key, A., D’Alessandro, A., and Palsson, B.O. (2025). RBC-GEM: A genome-scale metabolic model for systems biology of the human red blood cell. PLoS Comput Biol 21, e1012109.

103. Yao, H., Dahal, S., and Yang, L. (2023). Novel context-specific genome-scale modelling explores the potential of triacylglycerol production by Chlamydomonas reinhardtii. Microb Cell Fact 22, 13. 10.1186/s12934-022-02004-y.

104. Baldarelli, R.M., Smith, C.L., Ringwald, M., Richardson, J.E., and Bult, C.J. (2024). Mouse Genome Informatics: an integrated knowledgebase system for the laboratory mouse. Genetics 227. 10.1093/genetics/iyae031.

105. Cendali, F.A.-O., Lisk, C.A.-O., Dzieciatkowska, M., LaCroix, I.A.-O., Reisz, J.A.-O., Harral, J., Stephenson, D., Hay, A.A.-O., Wartchow, E.A.-O.X., Darehshouri, A.A.-O., et al. Increased exercise tolerance in humanized G6PD-deficient mice.

106. Ebrahim, A., Lerman, J.A., Palsson, B.O., and Hyduke, D.R. (2013). COBRApy: COnstraints-Based Reconstruction and Analysis for Python. BMC Syst Biol 7, 74. 10.1186/1752-0509-7-74.

107. Ebrahim, A., Beber, M.E., Mandal, S., König, M., Redestig, H., Diener, C., Scientist, D., John, P.S., akaviaLab, Hemant_Yadav, et al. (2024). opencobra/cobrapy: 0.29.1 (Zenodo).

108. Gudmundsson, S., and Thiele, I. (2010). Computationally efficient flux variability analysis. BMC Bioinformatics 11, 489. 10.1186/1471-2105-11-489.

109. Virtanen, P., Gommers, R., Oliphant, T.E., Haberland, M., Reddy, T., Cournapeau, D., Burovski, E., Peterson, P., Weckesser, W., Bright, J., et al. (2020). SciPy 1.0: fundamental algorithms for scientific computing in Python. Nat Methods 17, 261–272. 10.1038/s41592-019-0686-2.

110. Hunter, J.D. (2007). Matplotlib: A 2D graphics environment. Computing in science & engineering 9, 90–95.

111. Wu, T., Hornsby, M., Zhu, L., Yu, J.C., Shokat, K.M., and Gestwicki, J.E. (2023). Protocol for performing and optimizing differential scanning fluorimetry experiments. STAR Protocols 4, 102688. 10.1016/j.xpro.2023.102688.

112. Jumper, J., Evans, R., Pritzel, A., Green, T., Figurnov, M., Ronneberger, O., Tunyasuvunakool, K., Bates, R., Žídek, A., Potapenko, A., et al. (2021). Highly accurate protein structure prediction with AlphaFold. Nature 596, 583–589. 10.1038/s41586-021-03819-2.

