## Supplementary Figures 1-7 for "Structural divergence of G6PD variants shapes red blood cell storage and transfusion performance"

1) Department of Biochemistry and Molecular Genetics, University of Colorado Anschutz Medical Campus, Aurora, CO, USA;
2) Department of Pathology and Laboratory Medicine, University of North Carolina, Chapel Hill, NC, United States;
3) University of Virginia, Charlottesville, VA, USA;
4) RTI International, Research Triangle Park, NC, USA;
5) Omix Technologies Inc, Aurora, CO, USA;
6) Center for Biomedical Engineering and Technology, and Department of Physiology, University of Maryland School of Medicine, Baltimore, MD, United States of America.
7) Center for Blood Oxygen Transport and Hemostasis, Department of Pediatrics, University of Maryland, School of Medicine Baltimore, MD, United States of America.
8) Vitalant Research Institute, San Francisco CA, USA;
9) Department of Laboratory Medicine, University of California San Francisco, CA, USA
10) University of British Columbia, Victoria, British Columbia, Canada.
11) Structural Biology Initiative, CUNY Advanced Science Research Center, New York, NY, USA.
12) Biochemistry Krios Electron Microscopy Facility, Department of Biochemistry, University of Colorado Boulder, Boulder, CO, USA
13) Cryo-EM Structural Biology Shared Resource Facility, University of Colorado Anschutz Medical Campus, CO, USA
14) Shu Chien-Gene Lay Department of Bioengineering, University of California San Diego, La Jolla, CA, USA
15) Department of Pediatrics, University of California San Diego, La Jolla, CA, USA
16) Department of Pathology and Cell Biology, Columbia University Irving Medical Center, New York City, New York, USA.
17) Kaiser Permanente Northern California Division of Research, Oakland, CA
18) Department of Experimental and Clinical Medicine, University of Florence, Florence, Italy
19) Canadian Blood Services, Vancouver, Canada
20) Department of Pharmacology, University of Colorado Anschutz Medical Campus, CO, USA.
21) Department of Pharmaceutical Sciences, Skaggs School of Pharmacy, University of Colorado Anschutz Medical Campus, CO, USA.
22) BioFrontiers Institute, University of Colorado Boulder, CO, USA.
   Department of Pathology, University of Maryland, School of Medicine, Baltimore, MD, United States of America.
23) *These Authors Contributed Equally*
24) *Lead contact*

*** Correspondence:**
Angelo D'Alessandro, PhD
Department of Biochemistry and Molecular Genetics
University of Colorado Anschutz Medical Campus
12801 East 17th Ave., Aurora, CO 80045
Phone # 303-724-0096

www.dalessandrolab.com

John Janetzko
Department of Biochemistry and Molecular Genetics
University of Colorado Anschutz Medical Campus
12801 East 17th Ave., Aurora, CO 80045
 (J.J.)

SUPPLEMENTARY TABLE 1 ........ XSLX

### SUPPLEMENTARY FIGURES

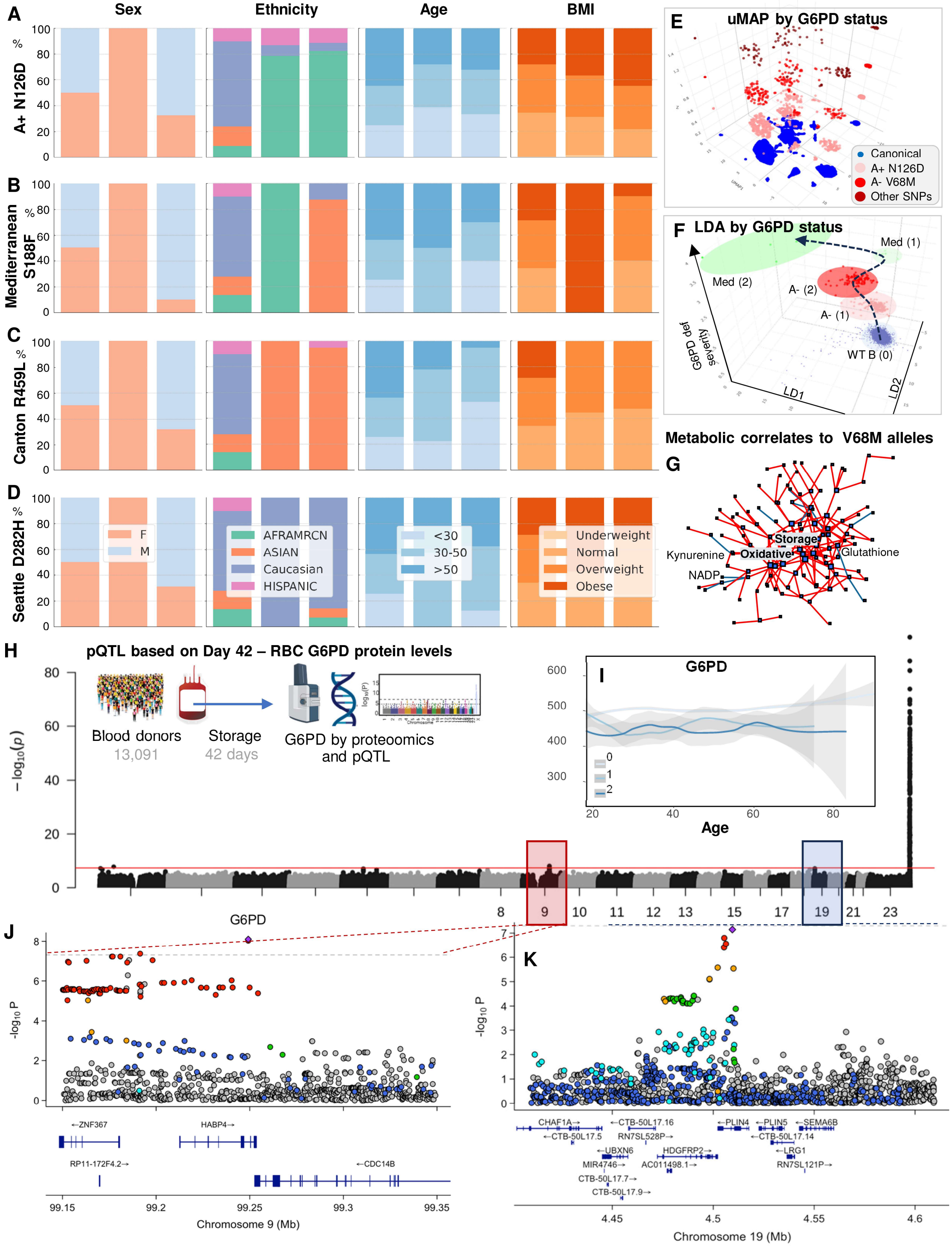

**Supplementary Figure 1 Demographic, metabolic, and protein-QTL features associated with G6PD variation in the REDS RBC-Omics index cohort.** Sex, self-reported ethnicity, age, and BMI distributions are shown for donors carrying N126D (A+) **(A)**, Mediterranean S188F **(B)**, Canton R459L **(C)**, and Seattle D282H **(D)**. UMAP of index-cohort RBC metabolomes is colored according to the predominant G6PD genotypes **(E)**.

Linear discriminant analysis of metabolomic profiles according to G6PD genotype illustrates separation of canonical, African A$^{-}$, and Mediterranean groups **(F)**. A network of metabolites associated with V68M allele dosage highlights redox-related metabolites and pathways **(G)**. Manhattan plot from the genome-wide pQTL analysis of day-42 RBC G6PD protein abundance is shown **(H)**. G6PD protein abundance is plotted as a function of donor age and genotype **(I)**. Locuszoom plots show the regional association of a major trans-pQTL signal on chromosome 9 **(J)** or 19 **(K)**.

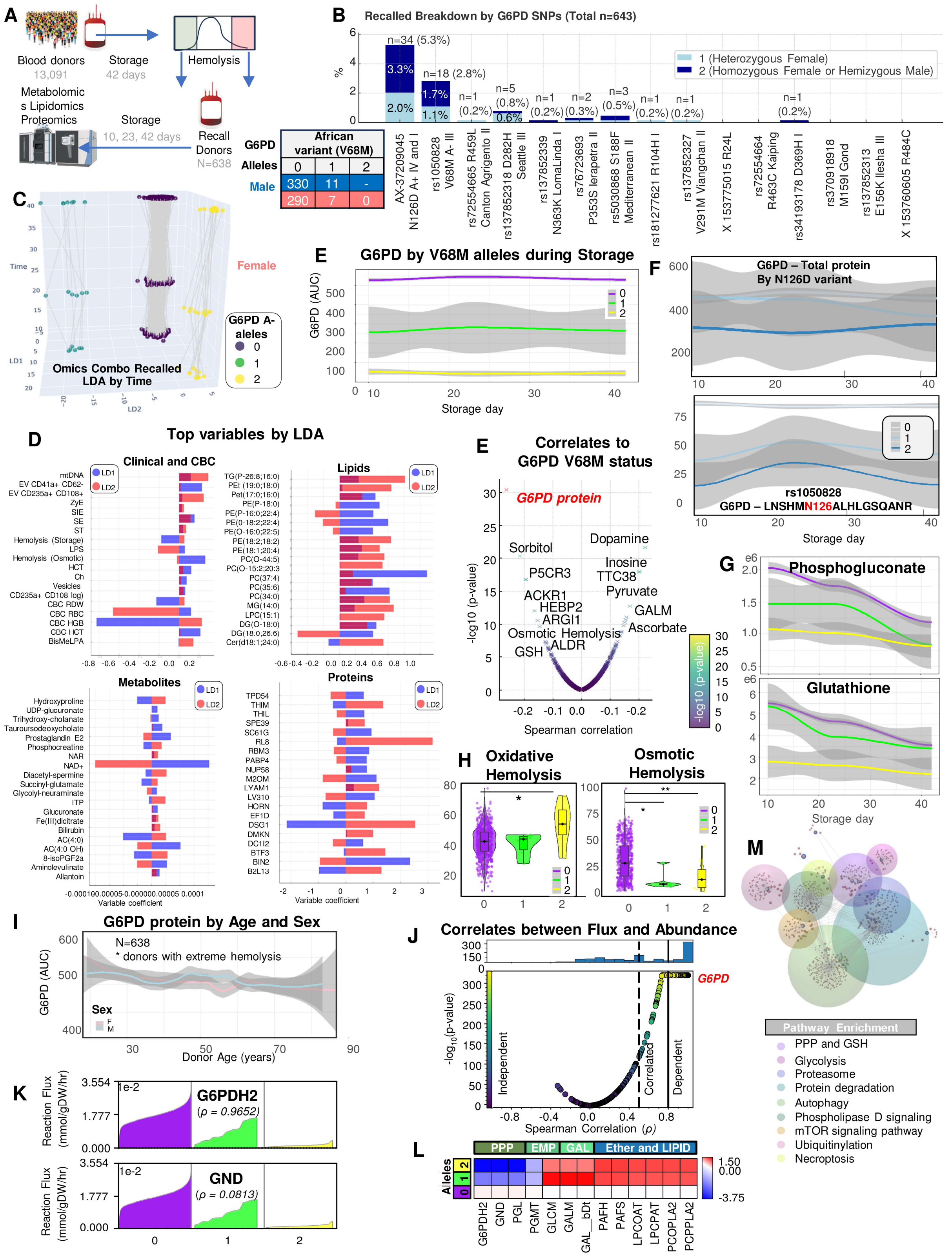

**Supplementary Figure 2 Recalled-donor multi-omics and proteome-constrained modeling validate G6PD allele-dose effects.** Overview of the REDS RBC-Omics recalled-donor study, in which 638 donors selected from extremes of storage, oxidative, or osmotic hemolysis donated a second RBC unit for longitudinal multi-omics analyses **(A)**. Frequencies of prevalent missense G6PD variants among recalled donors are shown, including the distribution of African A⁻ V68M allele dosage **(B)**. Three-dimensional linear discriminant analysis (LDA) of recalled-donor multi-omics data illustrates separation according to storage time and V68M allele dosage **(C)**. Top clinical/CBC, lipid, metabolite, and protein features contributing to discrimination according to V68M status are shown **(D)**. Multi-omics features correlated with V68M allele dosage are shown, with G6PD protein among the strongest negative correlates **(E)**. Longitudinal total G6PD protein abundance or selected peptide (including the N126 region) according to N126D or V68M allele dosage, together with representative PPP metabolites, is shown **(F-G)**. End-of-storage oxidative and osmotic hemolysis is shown according to V68M allele dosage **(H)**. G6PD protein abundance is plotted as a function of age and sex in recalled donors **(I)**. The relationship between modeled reaction flux and experimentally measured protein abundance classifies reactions as abundance-dependent, correlated, or independent **(J)**. Modeled effective flux ranges for representative significantly affected reactions, including PPP steps G6PD- (G6PDH2 in the model) and PGD (GND)-catalyzed reactions, are shown according to V68M allele dosage **(K)**. A heat map summarizes representative reactions with genotype-associated changes in modeled flux **(L)**. Pathway enrichment summarizes biological processes affected by G6PD genotype **(M)**.

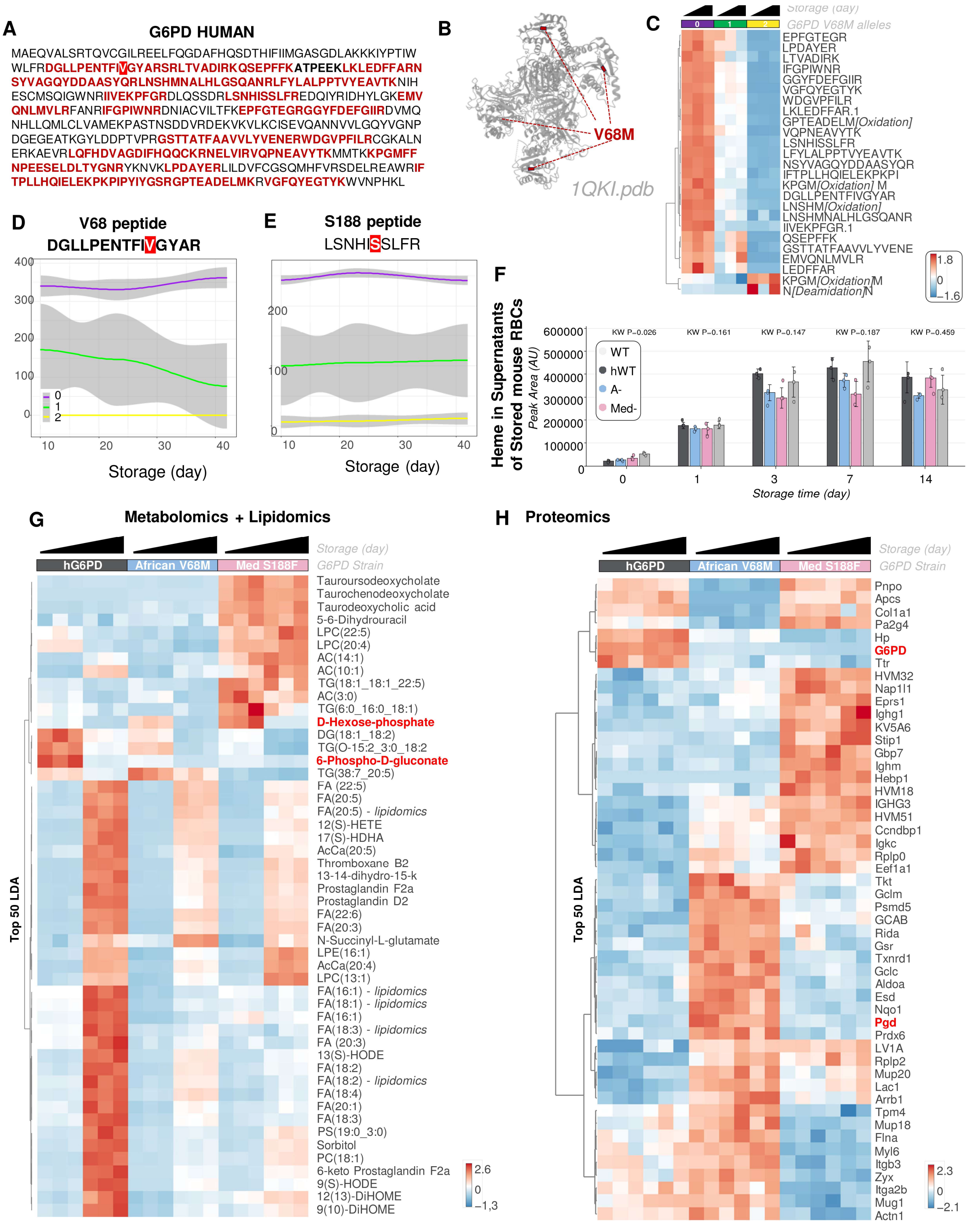

**Supplementary Figure 3 Peptide level data in REDS RBC Omics recalled donors and storage-dependent metabolomic, proteomic, and G6PD peptide signatures in humanized mouse RBCs.** Human G6PD protein sequence coverage in REDS RBC omics recalled donor samples **(A)**. V68M is mapped on the published human G6PD tetrameric structure of the Canton variant by Xray at 3Å (*1QKI.pdb*) **(B)**. Longitudinal abundance of the diagnostic V68-containing G6PD peptide is shown across storage duration (x axis) and genotype groups **(C)**. Longitudinal abundance of the V68M- and S188-containing diagnostic G6PD peptides is shown across storage and genotype groups **(D-E)**.

In the supernatants of stored murine RBCs from the WT C57BL/6J mice, or humanized mice expressing the canonical human G6PD or the A- and Med- variants, no significant changes were observed across strains with the respect to in bag hemolysis, as determined by free heme levels measured by mass spectrometry-based metabolomics **(F)**. A heat map shows the top metabolomic and lipidomic **(G)**, and proteomics **(H)** features distinguishing stored RBCs from humanized canonical, African A−, and Mediterranean G6PD mice across storage, including PPP, redox, and lipid-remodeling metabolites. In these humanized mice, mass spectrometry-based coverage of the human

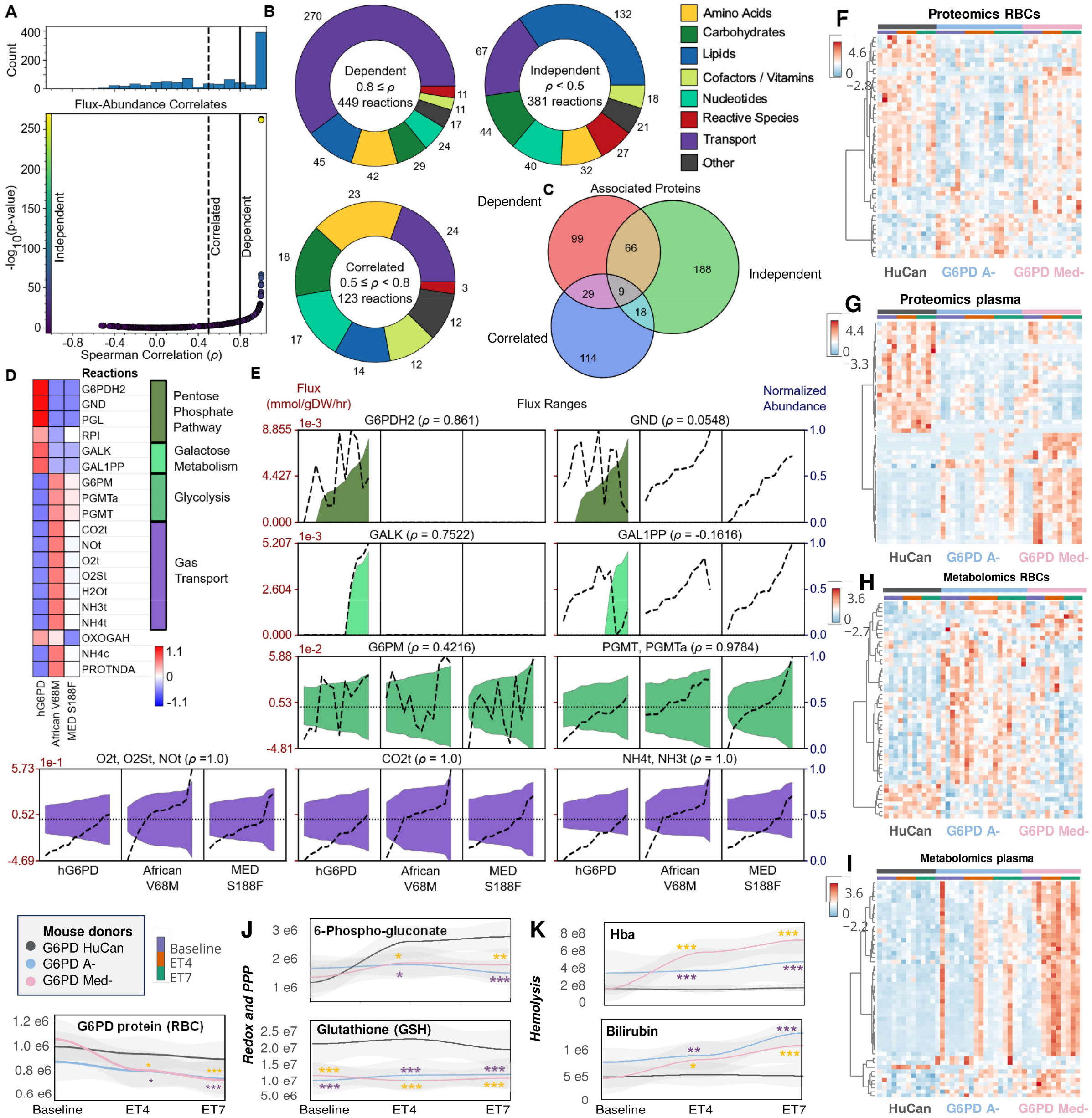

**Supplementary Figure 4. Systems-level metabolic modeling and multi-omics characterization of exchange transfusion with G6PD-deficient murine red blood cells.**

Relationship between experimentally measured protein abundance and predicted metabolic reaction flux across the proteome-constrained murine RBC metabolic model. Reactions were classified as abundance-dependent (Spearman $\rho \geq 0.8$), correlated ($0.5 \leq \rho < 0.8$), or abundance-independent ($\rho < 0.5$) **(A)**. Distribution of abundance-dependent, correlated, and independent reactions across metabolic pathway classes **(B)**. Overlap among proteins associated with abundance-dependent, correlated, and independent metabolic reactions **(C)**. Genotype-dependent differences in modeled flux ranges for representative reactions in the pentose phosphate pathway, galactose metabolism, glycolysis, and gas transport; dashed lines indicate normalized protein abundance and shaded areas indicate predicted flux ranges **(D–E)**. Heatmaps of RBC proteomic profiles following exchange transfusion with stored RBCs from humanized canonical G6PD (HuCan), African A−, or Mediterranean G6PD mice **(F)**. Heatmaps of recipient plasma proteomic profiles following exchange transfusion with the three donor RBC genotypes **(G)**. Heatmaps of recipient RBC metabolomic profiles following exchange transfusion **(H)**. Heatmaps of recipient plasma metabolomic profiles following exchange transfusion **(I)**. Longitudinal RBC G6PD protein abundance following exchange transfusion **(J)**. Longitudinal 6-phosphogluconate and reduced glutathione (GSH) levels following exchange transfusion, illustrating genotype-dependent alterations in pentose phosphate pathway and redox metabolism. Longitudinal plasma cell-free hemoglobin (Hba) and bilirubin following exchange transfusion, reflecting genotype-dependent differences in post-transfusion hemolysis **(K)**.

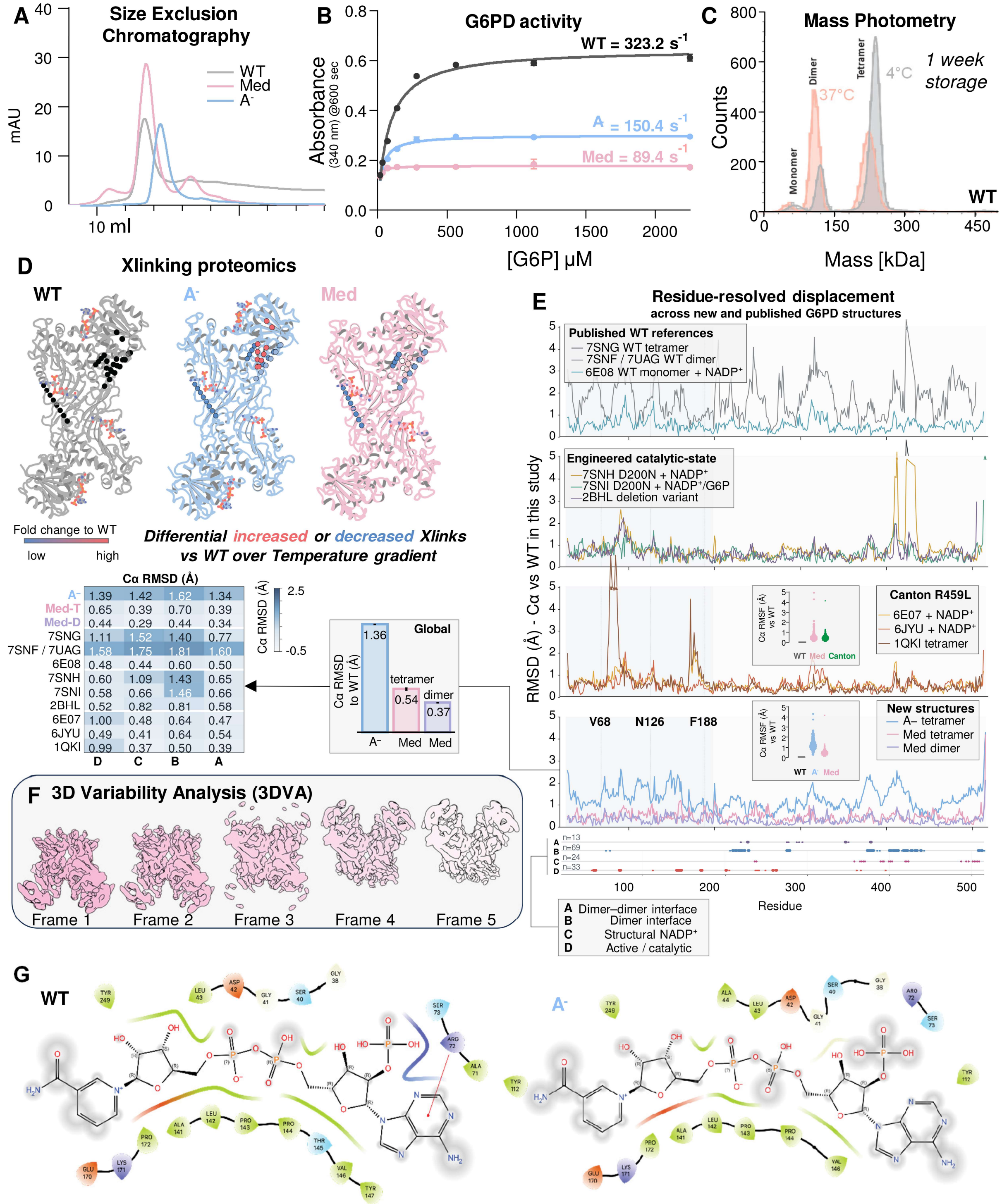

**Supplementary Figure 5. Additional biochemical, biophysical, and structural characterization of recombinant G6PD variants.** Size-exclusion chromatography profiles of recombinant WT, African A⁻ (V68M/N126D), and Mediterranean (S188F) G6PD **(A)**. Glucose 6-phosphate (G6P)-dependent NADPH production, measured by absorbance at 340 nm after 600 s. Lines show fitted substrate-response curves; annotated values indicate estimated catalytic turnover numbers (*k*cat, $s^{-1}$) **(B)**. Representative mass-photometry distributions of WT G6PD after one week at 4 or 37°C, with peaks corresponding to monomeric, dimeric, and tetrameric species **(C)**. TMT-based quantitative cross-linking proteomics across a temperature gradient, mapped onto WT, A⁻, and Mediterranean G6PD structures. Red and blue indicate increased and decreased cross-link abundance relative to WT, respectively, revealing variant-dependent changes in intramolecular and intersubunit contacts **(D)**. Residue-resolved Cα displacements following structural alignment to the NADP⁺-bound WT tetramer determined in this study. Comparisons include the newly resolved variants and published WT, engineered, and Canton R459L structures – as referenced in the manuscript. Bar plots and the heat map summarize global and functional-region Cα root-mean-square deviations (RMSDs), respectively; residue tracks identify the dimer–dimer interface, dimer interface, structural NADP⁺-binding region, and catalytic region **(E)**. Representative frames from three-dimensional variability analysis of the Mediterranean tetramer show coordinated conformational variability involving the region surrounding F188, consistent with heterogeneous local density in the consensus reconstruction **(F)**. Two-dimensional interaction maps compare the catalytic NADP⁺-binding environments of WT (left) and A− (right) G6PD **(G)**.

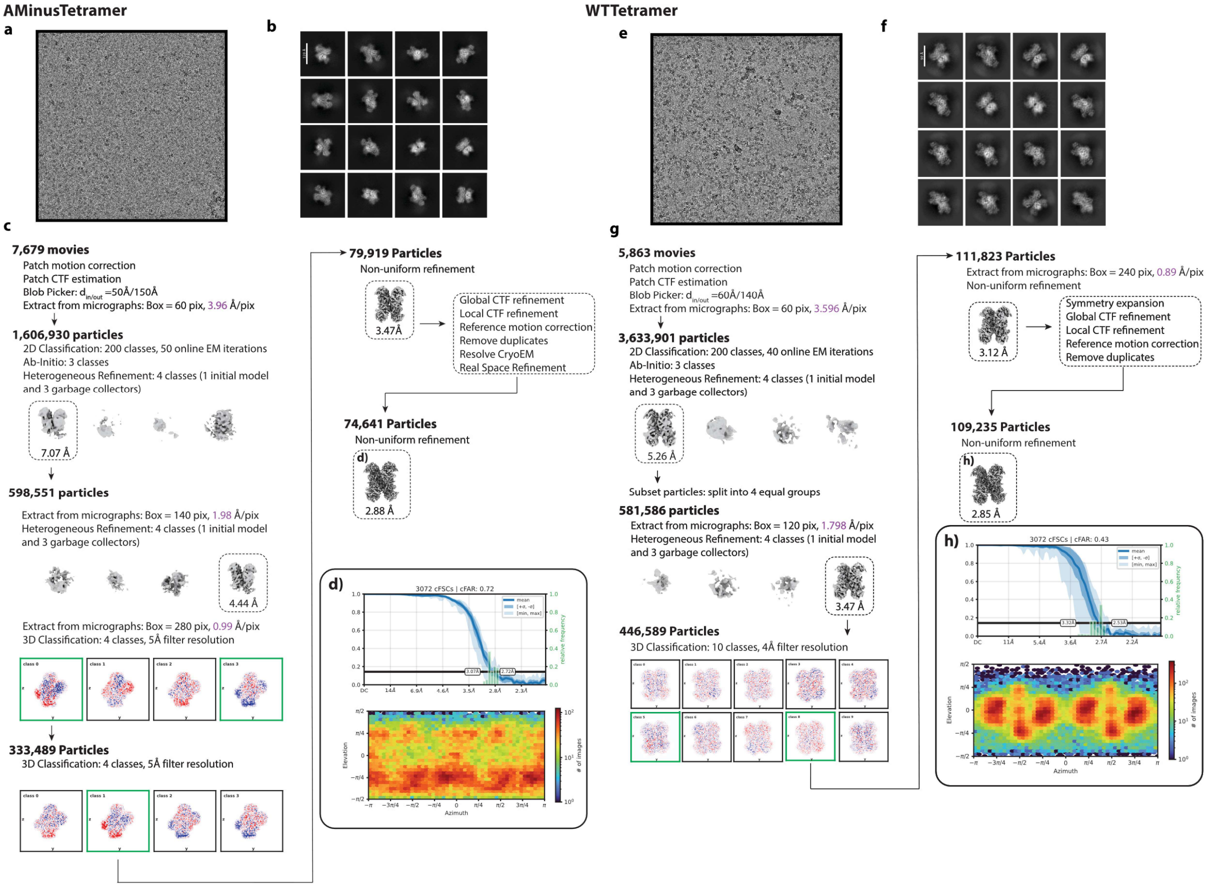

**Supplementary Figure 6 Cryo-EM processing and reconstruction of NADP⁺-bound WT, African A⁻, and Mediterranean G6PD oligomers.** Representative micrographs, two-dimensional class averages, particle-selection and classification workflows, three-dimensional refinements, Fourier shell correlation curves, and local-resolution distributions are shown for the African A⁻ tetramer and WT tetramer. Particle numbers, processing steps, map resolutions, and reconstruction parameters are indicated within the respective workflows. These analyses support the cryo-EM reconstructions presented in Figures 5 and 7.

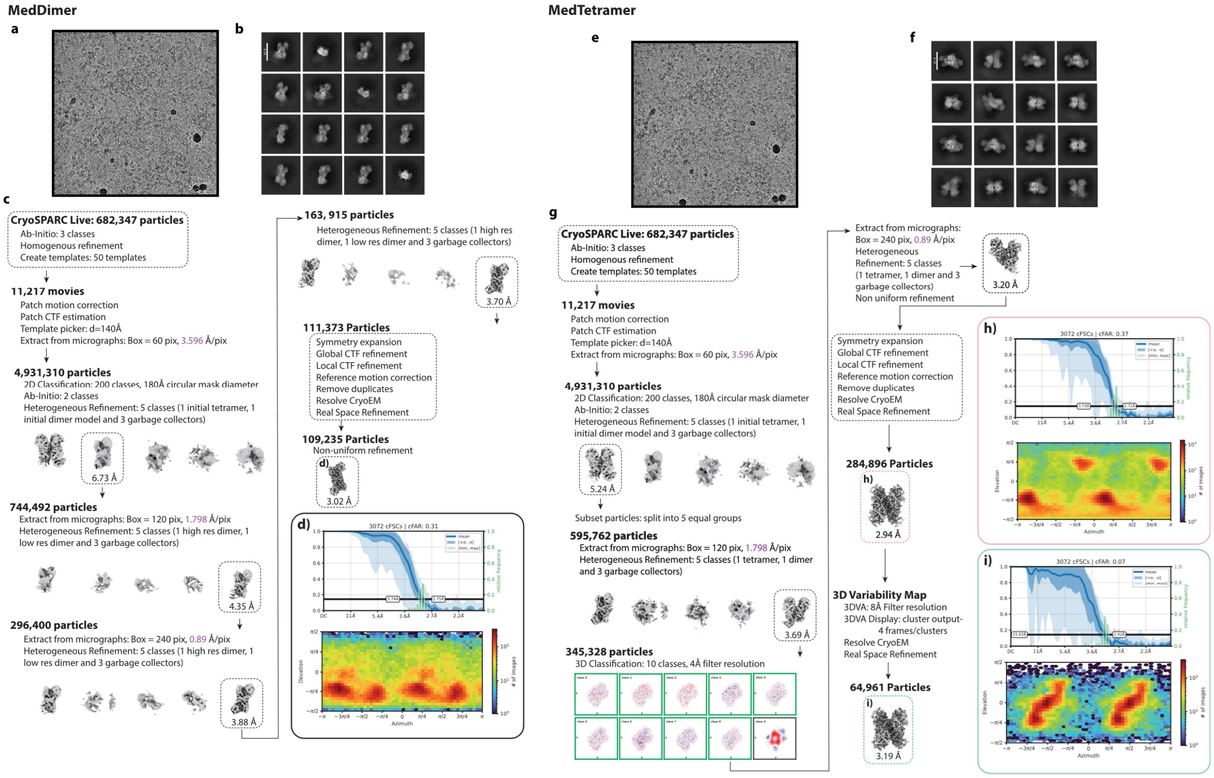

**Supplementary Figure 7 Cryo-EM processing and reconstruction of Mediterranean G6PD dimeric and tetrameric states.** Representative cryo-EM micrographs, two-dimensional class averages, particle-selection and classification workflows, three-dimensional refinements, Fourier shell correlation curves, and local-resolution distributions are shown for the Mediterranean S188F G6PD dimer and tetramer. Particle numbers, processing steps, map resolutions, and reconstruction parameters are indicated within the respective workflows. These analyses support the comparison of Mediterranean G6PD dimeric and tetrameric conformations presented in Figure 6.
